# Loss of Twist impairs tentacle development and induces epithelial neoplasia in the sea anemone *Nematostella vectensis*

**DOI:** 10.1101/2025.08.09.669484

**Authors:** Patricio Ferrer Murguia, Julia Hagauer, Emmanuel Haillot, Aissam Ikmi, Alison G Cole, Ulrich Technau

**Affiliations:** Dept. of Neurosciences and Developmental Biology, Faculty of Life Sciences, University of Vienna, Djerassiplatz 1, 1030, Vienna; Developmental Biology Unit, European Molecular Biology Laboratory, 69117 Heidelberg, Germany; Research platform “Single cell regulation of stem cells”, University of Vienna, Djerassiplatz 1, 1030 Vienna, Austria; Max Perutz labs, University of Vienna, Dr. Bohrgasse 5, 1030 Vienna, Austria

**Author notes:** Friedrich Miescher Laboratory of the Max Planck Society, Max-Planck-Ring 9,72076, Tübingen, Germany (Julia Hagauer).

## Abstract

The basic helix-loop helix transcription factor Twist plays diverse roles in mesodermal development across bilaterians, but its function in cnidarians remains unclear. Here, we investigate the role of Twist in tentacle morphogenesis and tissue homeostasis in the sea anemone *Nematostella vectensis*. Using a CRISPR/Cas9 generated knockout, we show that *twist* mutants exhibit impaired secondary tentacle formation, reduced proliferation in budding tentacles. Cross-sections reveal that mutants also lack micronemes, which are incomplete mesenteries that demarcate tentacle boundaries-suggesting defects in spatial patterning. We demonstrate that *twist* expression is regulated by Wnt, BMP, and Notch signalling but is independent of MAPK and Hedgehog pathways. Loss of Twist disrupts expression of mesodermal transcription factors *paraxis* and *tbx15* and perturbs the TOR-FGF signalling feedback loop necessary for normal tentacle growth. In addition to the impaired tentacle formation phenotype, juvenile or adult mutants develop epithelial neoplasms at the level of the pharynx, with tentacle-like molecular and morphological profiles, indicating a role for Twist in maintaining tissue homeostasis at the oral pole. Together, our findings reveal that Twist integrates major signalling pathways to regulate secondary tentacle patterning and maintain spatial tissue organisation in the diploblastic *Nematostella vectensis*.

## Introduction

Appendages, such as legs, fins, antennae and tentacles are common morphological structures in many animals. Appendages have many important roles such as locomotion, feeding, and sensory perception (Abzhanov et al., 2001; Fritz et al., 2013; Heming, 1996; Panganiban et al., 1997). In Cnidaria, these outgrowths form the tentacles and are mostly used to catch prey, sense the environment and act in defence. They can vary in morphology and number, but all share the common feature of having a high density of stinging cells, the phylum-defining cell types (Fritz et al., 2013; Fujiki et al., 2019; Gold et al., 2015; Zenkert et al., 2011). What controls the formation of the tentacles in a highly defined spatio-temporal pattern, remains largely unsolved.

In the sea anemone *Nematostella vectensis*, a well-established cnidarian model organism (Genikhovich and Technau, 2009a; Hand and Uhlinger, 1992), the 16 tentacles form at the oral pole as extensions of the body wall, surrounding the mouth. Four tentacle anlagen are formed already during larval development, and they form the first four tentacles present in the primary polyp. The polyp has eight mesenteries that form during larval development. They divide the animal into eight segments, which serve as initial boundaries between neighbouring tentacles (He et al., 2018; Ryan et al., 2007). The four primary tentacles of the primary polyp are positioned in every other segment, defined by the mesenteries. The onset of tentacle development and extension is marked by changes in the ectoderm forming a placode, followed by modifications in the epithelial cell layer that drive elongation (Fritz et al., 2013). The next four tentacles form in the segments between the primary tentacles. Later, in a feeding-dependent manner and through several stereotyped phases, the other secondary tentacles are added, commonly resulting in a total of sixteen tentacles (Ikmi et al., 2020). To achieve this, each segment must be divided through the formation of micronemes, which appear as incomplete mesenteries and act to delineate boundaries between neighbouring tentacles.

Some of the molecular mechanisms for normal tentacle development in *N. vectensis* have already been deciphered. Specifically, in developing larvae, a set of Hox genes, together with *gbx*, are expressed in the gastric tissue of the planula larva in a staggered manner along the directive axis, which is perpendicular to the main oral-aboral axis. Their expression boundaries mark precisely the position of the later mesenteries (He et al., 2018). All hox genes and *gbx* are regulated by a BMP signalling gradient along the directive axis (Genikhovich et al., 2015; Saina et al., 2009) and delineate the segments necessary for initial tentacle development and patterning (Finnerty et al., 2004; He et al., 2018; Ryan et al., 2007).

In a recent study, FGF signalling was found to play a role in tentacle formation. Specifically, a crosstalk between Fibroblast Growth Factor Receptor b (FGFRb) and Target of Rapamycin (TOR) in the gastrodermal layer drives the development of secondary tentacle growth (Ikmi et al., 2020). At present it is unknown what regulates the local activation of the FGFR expression in the early tentacle bud anlagen. In some bilaterian species, the bHLH transcription factor Twist acts upstream of FGFR signalling, regulating various developmental processes, including the growth and extension of appendages (Hornik et al., 2004; Lu et al., 2012; Zhu et al., 2014). Notably,in *Nematostella, twist* is expressed in the mesodermal lining of the pharynx and at lower levels in the body wall (Martindale et al., 2004), the region, where TOR signalling activates FGFR signalling during secondary tentacle formation. However, the upstream regulation of *twist* and its role in tentacle morphogenesis in *Nematostella* remain unexplored.

Here, we report that *twist* is required for proper formation of secondary tentacles through regulation of the TOR signalling pathway. We further show that *twist* expression is regulated by Wnt, Notch, and BMP signalling, while MAPK and Hedgehog inhibition do not affect its expression. Moreover, in juvenile to subadult stages of the polyp, *twist* mutants form amorphic hyperplasia of the body wall with tentacle tissue identity, suggesting that Twist might be necessary for localized and directed outgrowth of the tentacle appendages. These results indicate that Twist functions at the intersection of major developmental signalling pathways and a mesodermal-like transcriptional program in *Nematostella*.

## Results

### *twist^−/−^* mutants develop 8 mesenteries and shorter primary tentacles

Expression analyses by in situ hybridisation confirmed that *twist* expression is not detectable during early gastrulation (Fig. 1A), and only faintly expressed in the 2-day-old planula (Fig. 1B). By 72 hours post-fertilization expression becomes clearly detectable in the inner layer of the pharynx (Martindale et al., 2004; Fig. 1C). Expression is maintained throughout the late planula and in the polyp stage (Fig. 1D,E). Notably, in the polyps, the expression domain appears split into two distinct regions; one at the outer pharynx below the tentacles and one in the lower, inner pharynx (Fig. 1E). As the animal transitions into the polyp, *twist* expression adopts a “cage“-like pattern around the pharynx being more prominent at the bottom of the pharynx and around of the primary tentacles.

**Figure 1:**
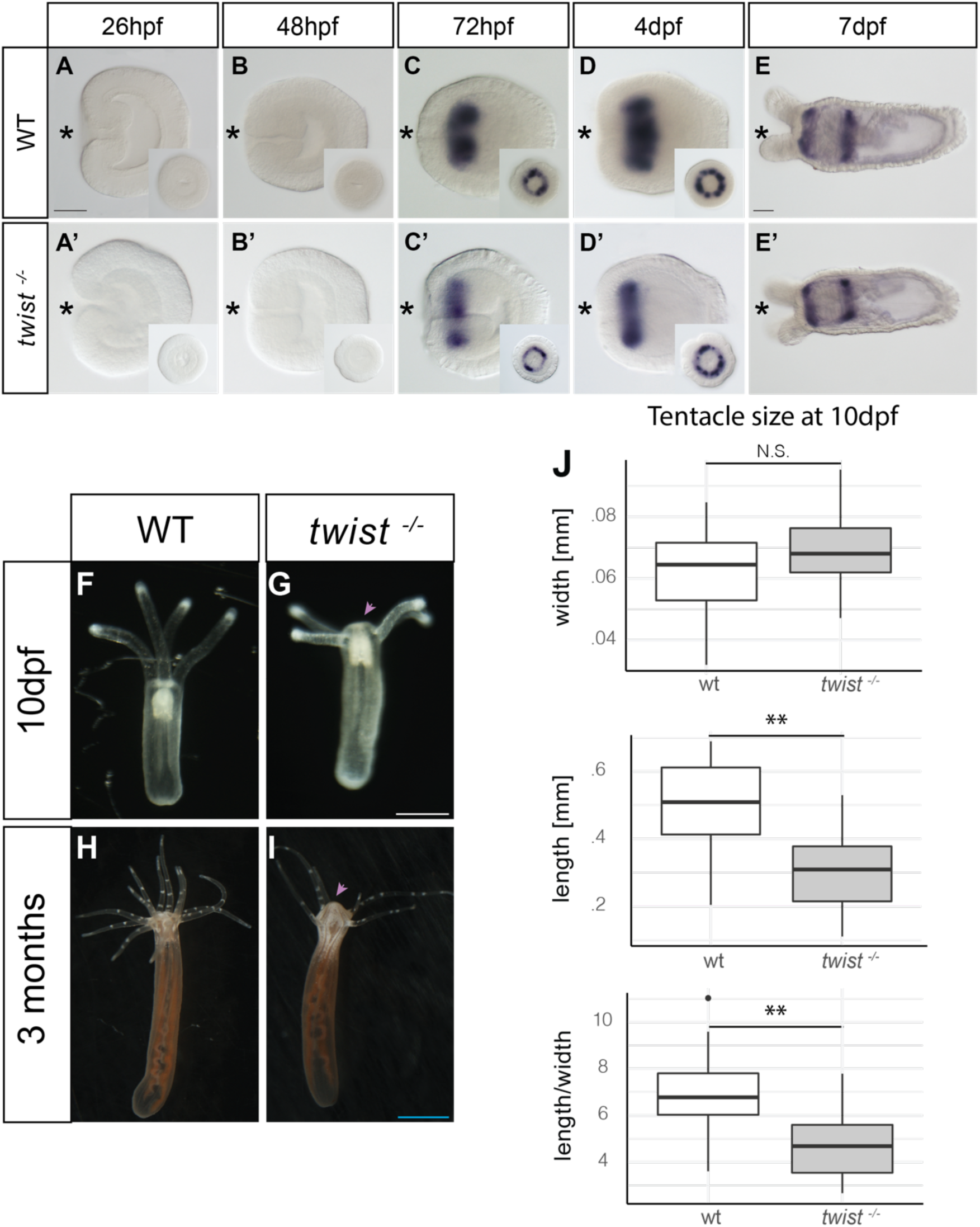
*twist^−/−^* early mutant phenotype in *N. vectensis.* Whole Mount In Situ Hybridization (WMISH) *twist* expression in *Nematostella vectensis* at different developmental stages, **(A. A’)**26h late gastrula, (B, C, B’, C’)48h-4dpf planulae and **(E, E’)**7dpf primary polyp, comparing the WT and *twist^−/−^* mutants. Dark blue indicates expression. Asterix indicates oral pole of the animals. **(F)** Wild-type (WT) primary polyp at 10 days post-fertilization (dpf). **(G)** *twist^−/−^*mutant polyp at the same age as in (A). The oral disc of *twist^−/−^*animals exhibits a protrusion (pink dashed line arrowhead) instead of flat mouth as in wildtype (white dashed line). **(H)** WT polyp at 3 months of age, illustrating normal development. **(I)** *twist^−/−^* mutant polyp, showing the phenotype characterized by altered oral disc morphology (Dashed lines). **(J)** Boxplot displaying the width (left), length (right), and ratio (bottom) of tentacle measurements in WT and *twist* mutant animals. At 10 dpf, primary tentacles in WT animals are longer compared to *twist^−/−^* mutants, as indicated by the boxplot analysis. Scale bars are 50µm, Black, 500µm White, 5mm Blue.

To study the function of the TF Twist in the sea anemone *Nematostella vectensis* we generated a *twist* knockout using CRISPR/Cas9 (for details see Materials and Methods). The 5 bp deletion induces a frameshift after the first helix, introducing a premature stop codon after 67 amino acids. The resulting truncated protein lacks the loop and second helix of the bHLH domain and is thus unable to form dimers (Sup. Fig. 1A-C). Since bHLH proteins form obligatory dimers, this mutation is predicted to be non-functional in transcriptional activation. Interestingly, the homozygous *twist* mutant still shows the same expression pattern as the wildtype, suggesting that nonsense-mediated decay may not be active in *Nematostella* (Fig. 1A’-E’).

Although early expression of *twist* is detected in the planula stage, no changes in the size or development of the eight pre-mesenterial segments were observed in the mutants. (Sup. Fig. 2 & Mov.1,2). Notably, at 10 days post-fertilization (dpf), morphological differences between WT and *twist^−/−^* animals become apparent. The oral disc of *twist^−/−^* animals displays a protrusion, which becomes increasingly pronounced as animals mature, somewhat reminiscent of the hypostome of hydrozoans (Fig 1G,I). Additionally, the primary tentacles in WT animals are somewhat longer than those of *twist^−/−^* mutants at the same developmental stage (Fig. 1J).

### Twist expression is regulated by Wnt, Notch and BMP signalling and required for mesodermal transcription networks in *Nematostella*

To determine how *twist* expression is regulated, we pharmacologically inhibited key signalling pathways (predominantly 48-72 hpf) and analysed *twist* expression by in situ hybridization (Fig. 2A, B). Our results reveal that *twist* expression is regulated by Wnt, BMP and Notch signalling but is independent of MAPK and Hedgehog signalling. Ectopic activation of the Wnt/ß-catenin pathway led to an expansion of *twist* expression throughout the whole mesoderm (Fig. 2A). Conversely, inhibition of Wnt signalling completely abolished *twist* expression (Fig. 2A). Additionally, both Notch and BMP signalling are required for normal *twist* expression. Notch inhibition resulted in a total loss of *twist* expression while BMP signalling inhibition significantly reduced or abolished *twist* (Fig 2B). In contrast, inhibition of MAPK or Hedgehog signalling had no effect, indicating that these pathways do not regulate *twist* expression in *Nematostella* (Fig. 2B).

**Figure 2.**
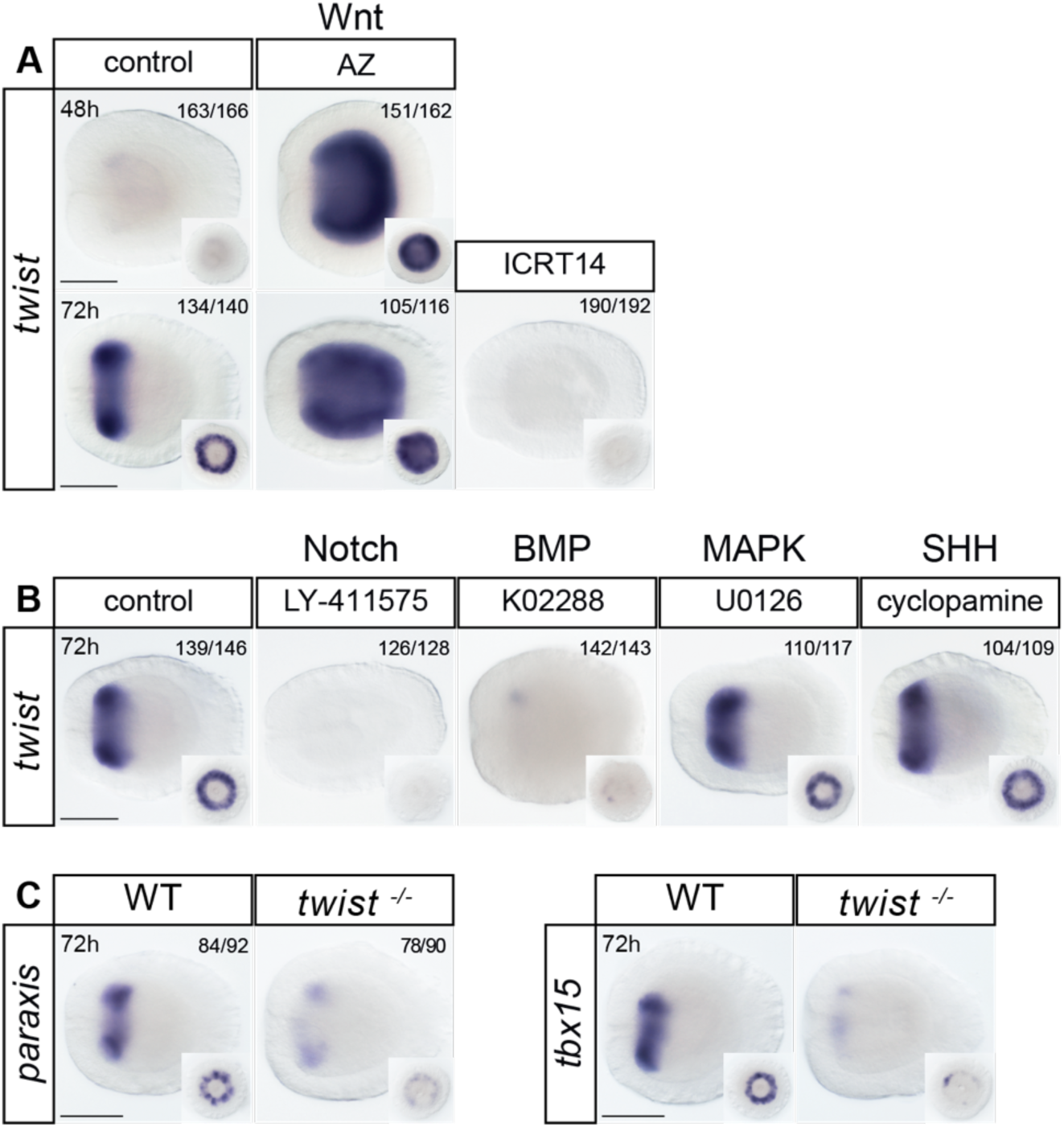
*twist* is regulated by Wnt, Notch and BMP signalling and required for *paraxis* and *tbx15* expression. **(A)** *twist* expression at 48 h and 72 h following modulation of Wnt signalling. Activation with azakenpaullone (AZ) expands *twist* expression, while inhibition with ICR14 abolishes expression. **(B)** Inhibition of Notch with LY-411575 and of BMP with K02288 reduces or eliminates *twist* expression. Inhibition of MAPK with U0126 or of SHH with cyclopamine has no detectable effect. **(C)** Expression of *paraxis* and *tbx15* is reduced in *twist*–/– embryos compared with wild type. Lateral views; oral views shown as insets. Scale bars: 50 µm. Controls were all in DMSO.

To further explore the molecular consequences of the complete loss of Twist, we examined the expression of two mesodermally co-expressed TFs, *paraxis* and *tbx15* in the *twist^−/−^* mutants. Both TFs showed strong downregulation in the mutant when compared to the WT (Fig. 2C). This indicates that Twist is upstream and necessary for the expression and/or maintenance of *paraxis* and *tbx15*, placing *twist* upstream within the pharyngeal mesodermal transcriptional regulatory network in *Nematostella*.

### Secondary tentacle formation is severely delayed or abolished in the *twist* mutant

At roughly 14–15dpf, development of secondary tentacle begins, which is promoted by feeding (Ikmi et al., 2020). To ascertain whether additional tentacle development was hampered, we started to feed animals 10dpf every day. To make sure that *twist* mutants and WT were able to feed properly, the animals’ growth (length and width) was also monitored. No initial differences in body length were observed, only after 13 days of constant feeding did the *twist^−/−^* animals become significantly shorter than the WT polyps, with an average reduction length of 15.5%. While the WT polyps are initially slightly narrower than the mutant polyps, no statistically significant difference is observed in later measurements. The body length to width ratio of the mutant is considerably lower at all recorded time intervals, showing that mutant polyps of the same length frequently have broader mouth regions (Fig. 3A). The most striking phenotype of the *twist^−/−^*mutants at the juvenile polyp stage is found in the formation of secondary tentacles. Mutants mostly only developed 4-8, at maximum 10 tentacles, as opposed to the WT’s typical 16 tentacles, and secondary tentacle addition was significantly delayed in *twist^−/−^* mutants compared to WT animals (Fig. 3B). Some mutants retained only the four primary tentacles, without ever fully developing secondary tentacles (Fig. 3b’). Moreover, if secondary tentacles were formed, they were significantly thinner and shorter in *twist^−/−^* mutants (Fig. 3b,b’; Sup. Fig 3). Because later phases of secondary tentacle addition are typically followed by pseudo-segmentation of the eight body chambers, so called micronemes, we looked for these mesodermal folds in cross-sections of polyps in adult polyps. Interestingly, while the micronemes were present in all WT animals, they were completely absent in the mutants (Fig. 3C).

**Figure 3.**
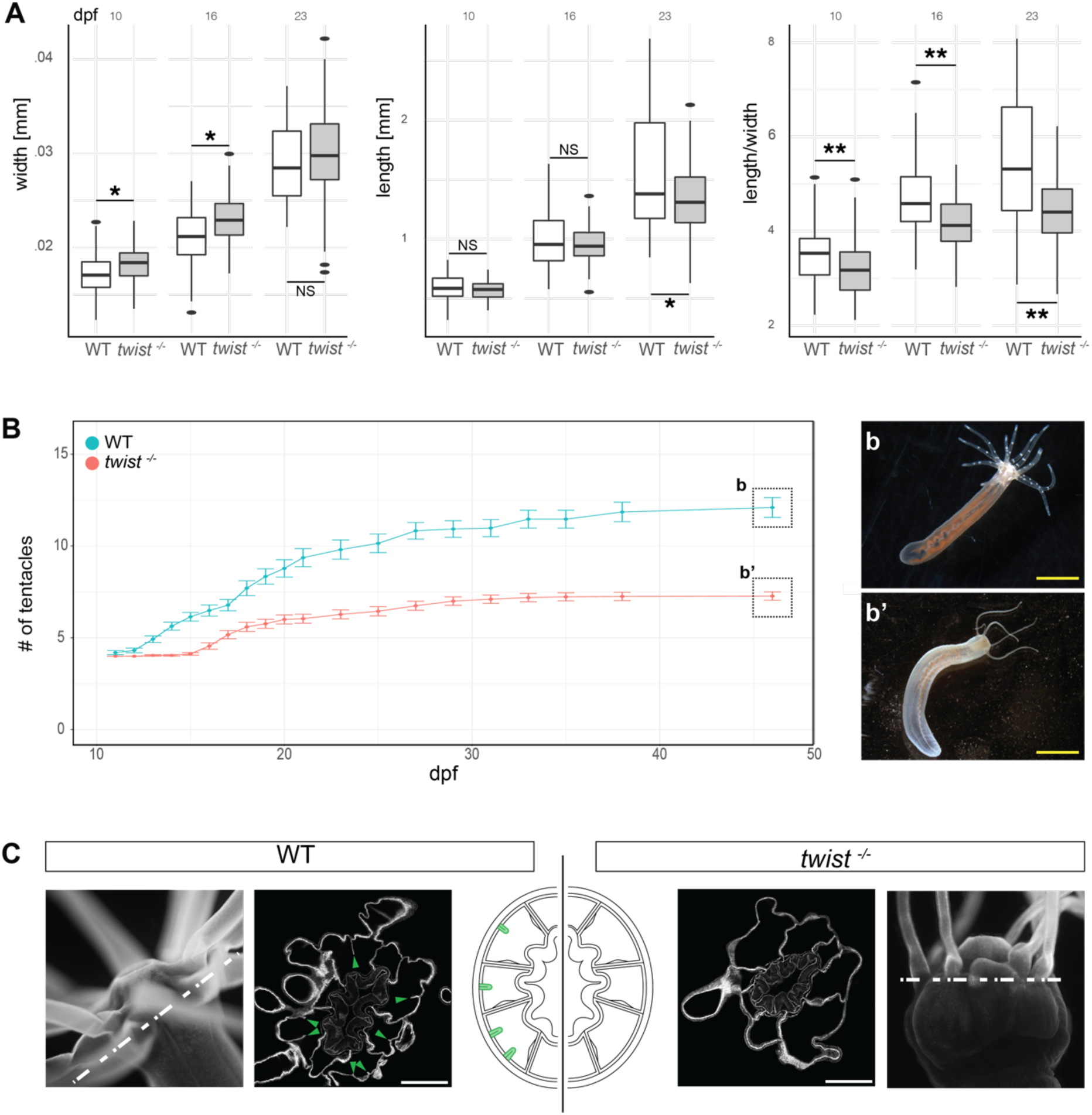
Twist mutants show altered body proportions, impaired tentacle development, and lack oral micronemes. **(A)** Quantification of body width (left), length (center), and width-to-length ratio (right) in wild-type (WT) and *twist^−/−^* polyps. **(B)** Tentacle number in WT (cyan) and *twist^−/−^* (red) animals across 46 days post-fertilization (dpf). **(b)** WT polyp at 46 dpf with 12 fully formed tentacles. **(b’)** *twist^−/−^* mutant at the same age with 4 fully developed tentacles. **(C)** Confocal cross-sections at the oral pole stained with phalloidin (grey). WT animals show distinct oral micronemes (green arrowheads), absent in *twist^−/−^* mutants. A schematic (middle) illustrates oral pole organization in WT and *twist^−/−^*animals, highlighting the presence or absence of oral micronemes at the base of the body wall. Scale bars 100 µm white, 50mm yellow. Dashed line indicates a sagittal section.

In wildtype animals, the outgrowth of new secondary tentacles is preceded by local spots of proliferation (Ikmi et al., 2020). We therefore performed EdU pulse labelling on growing tentacle buds. While the early and elongating tentacle buds of WT polyps showed localised proliferation as reported before (Ikmi et al., 2020), the early buds of mutant polyps did not exhibit any increased proliferation although they still display the morphological formation of a bud (Fig. 4A,B). This means that the thickening of the early bud tissue is independent of cell proliferation. We next decided to investigate the TOR and FGF pathways in the *twist^−/−^* mutants. We used immunostaining to detect the presence of phosphorylated ribosomal protein S6 (pRPS6), a marker of TOR activity that has been shown to play a role between nutrient uptake and signalling processes in the development of secondary tentacles (Ikmi *et al*., 2020). In contrast to WT animals, in which pRPS6 is detected in both initial and extending tentacle buds, pRPS6 was exclusively detected in extending tentacle buds in the *twist^−/−^*mutant (Fig. 4C,D).

**Figure 4:**
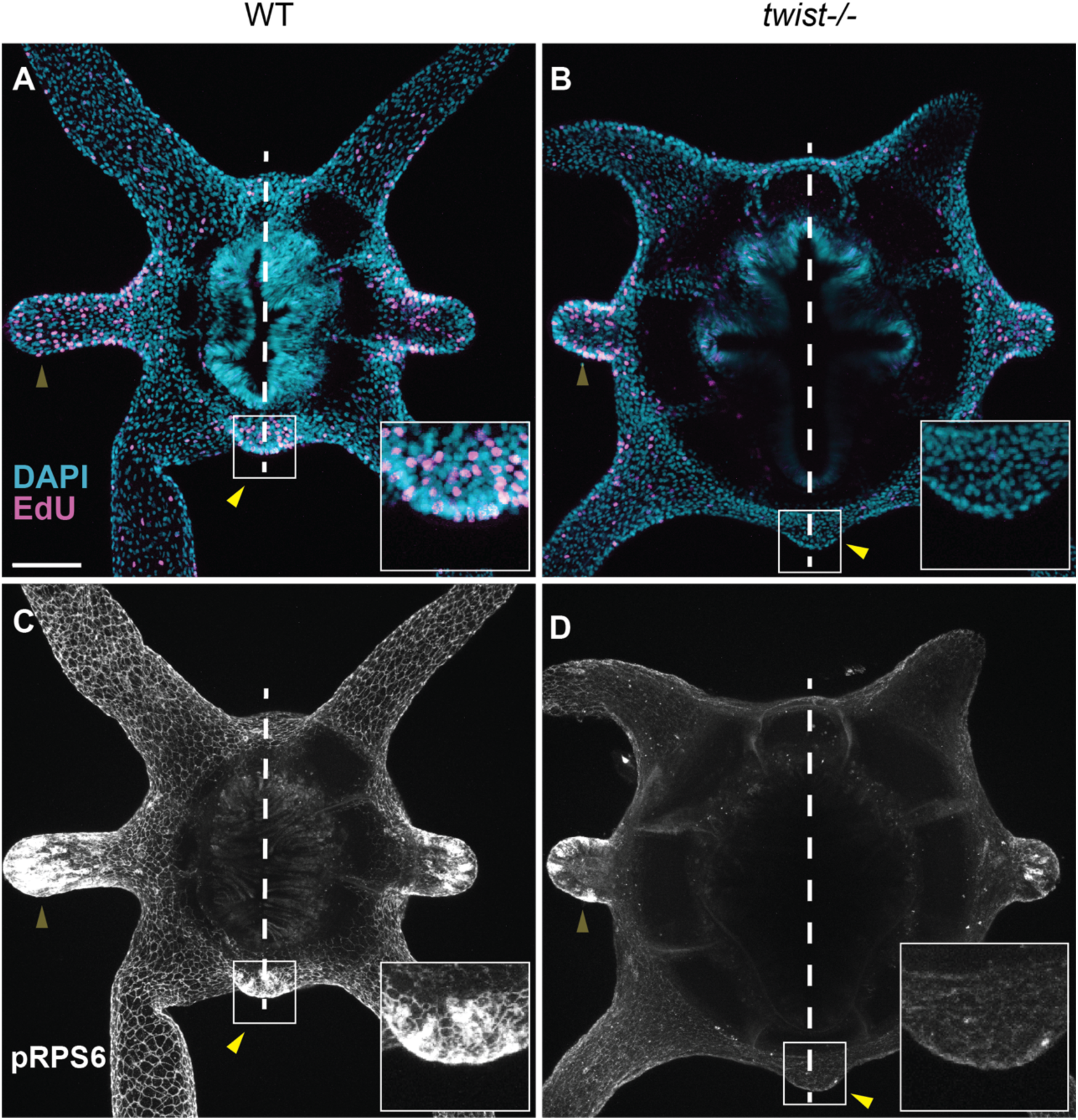
Tentacle bud development and TOR Activity is impaired in *twist* mutants. EdU staining of an overhead view of WT polyp. **(B)** EdU staining of an overhead view of twist mutant polyp. **(C)** Immunostaining for pRPS6, indicating TOR pathway activity, in WT polyp. **(D)** Immunostaining for pRPS6 in twist mutant polyp. Yellow arrowheads indicate early tentacle buds, while light yellow arrowheads indicate secondary tentacles that are already elongating. Dashed lines are used to indicate the directive axis. Scale bar 100 μm.

The spot-like expression domain of FGFRb expression at the site of the future tentacle outgrowth could be visualized in a FGFRb::eGFP reporter line (Ikmi et al., 2020). To explore the role of Twist in secondary tentacle development, we generated a *twist^−/−^* / Fgfrb::eGFP mutant transgenic line to examine any changes in the Fgrfb-positive ring muscle cells in both fed and unfed polyps. In unfed polyps, both WT and *twist^−/−^* mutants initially exhibited similar levels of detectable Fgfrb::eGFP positive cells (Fig. 5A’,F’). However, once the animals were fed and began to mature, the Fgfrb::eGFP positive cells in *twist^−/−^* mutants became weaker and smaller, though not completely abolished (Fig. 5B’,G’). Furthermore, as the secondary tentacles started to elongate, the Fgfrb::eGFP positive cells in *twist^−/−^* mutants became significantly weaker compared to the WT (Fig. 5D’,I’). Our observations suggest that Twist plays an important role in maintaining the signalling dynamics between the TOR and FGF pathways, which are critical for localized proliferation and proper tentacle growth.

**Figure 5.**
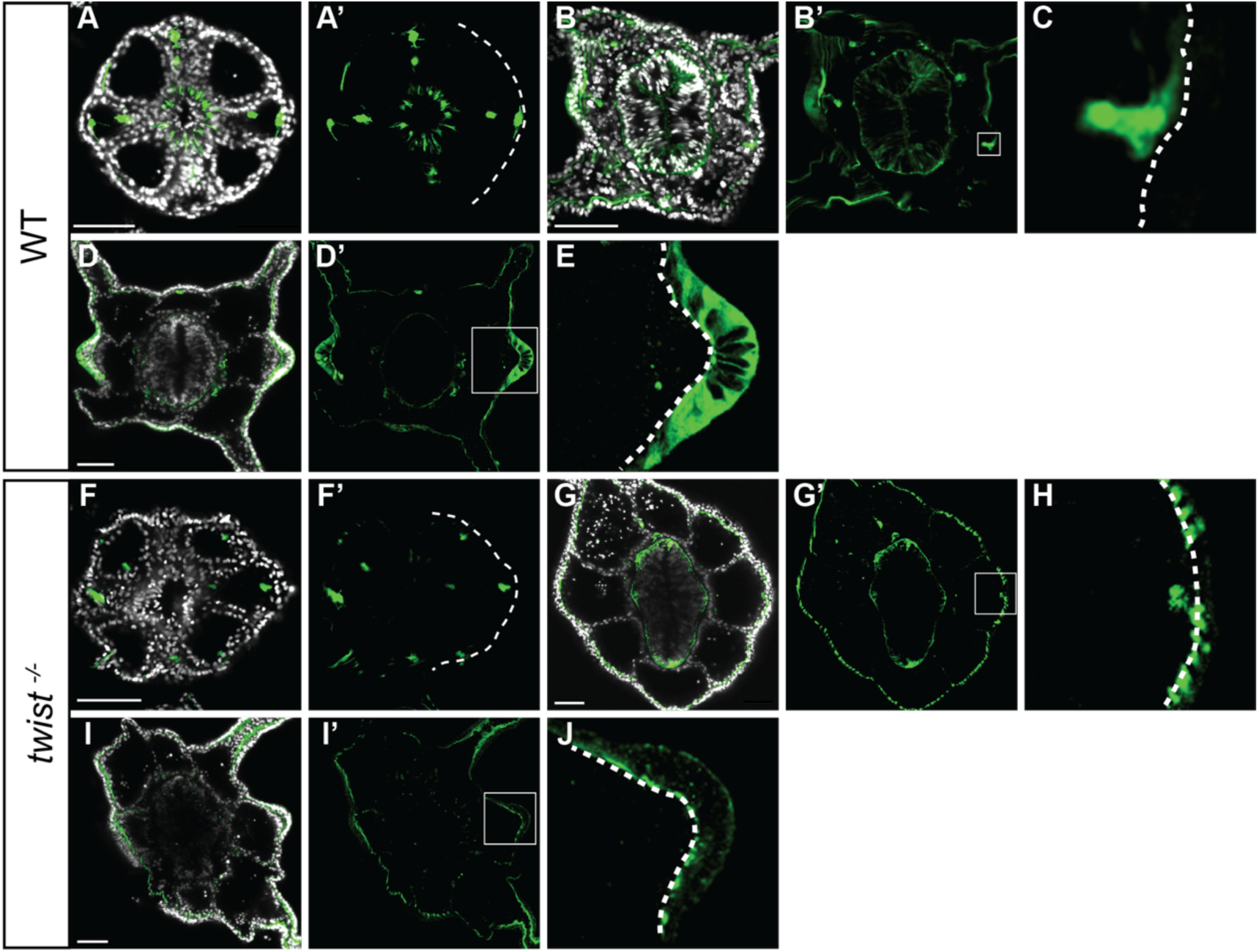
*Fgfrb-eGFP* expression in budding tentacles is reduced in *twist ^−/−^* animals. **(A,A’)** Expression of *Fgfrb-eGFP* in ring muscles of non-fed WT animals, with DAPI staining in (A). **(B,B’)** Expression of Fgfrb-eGFP in WT animals 4 days after feeding, with DAPI staining in (B). **(C)** Close-up view of *Fgfrb-eGFP* expression in ring muscles of WT animals 4 days after feeding. **(D,D’)** Expression of Fgfrb-eGFP in secondary WT tentacle buds, with DAPI staining in (D). **(E)** Close-up view of Fgfrb-eGFP expression in secondary WT tentacle buds. **(F,F’)** Expression of Fgfrb-eGFP in ring muscles of non-fed *twist^−/−^* animals, with DAPI staining in (F). **(G,G’)** Expression of Fgfrb-eGFP in *twist-/-* animals 4 days after feeding, showing diminished expression, with DAPI staining in (G). **(H)** Close-up view of diminished *Fgfrb-eGFP* expression in ring muscles and initial tentacle buds of *twist^−/−^* animals 4 days after feeding. **(I, I’)** Expression of *Fgfrb-eGFP* in secondary *twist^−/−^* tentacle buds, with DAPI staining in (I). **(J)** Close-up view of *Fgfrb-eGFP* expression in secondary *twist^−/−^* tentacle buds, showing significantly reduced signal. Scalebar100 μm

Since secondary tentacles in the *twist^−/−^* mutants do not develop normally, we wondered whether the structure and composition of the tentacles is normal. First, we assessed the tentacle retractor muscle. Most muscles in *Nematostella* are located in the gastrodermis, with the exception of the tentacle retractor muscles, which are of ectodermal origin (Jahnel et al., 2014; Cole et al., 2023). The tentacle ectodermal retractor muscles are longitudinally orientated, whereas gastrodermal muscles are circular. We find that muscle organisation is maintained even if secondary tentacles in *twist^−/−^* mutants are shorter and often thinner than those found in WT animals (Sup. Fig 4A,A’,B,B’). We conclude that *twist* has no role in muscle formation of the tentacles, as in the rest of the body. Secondly, we checked whether the three cnidocyte types (basitrichous haplonema, microbasic mastigophore, and spirocytes) that are typically found in WT tentacles, were also present in the mutant tentacles. Late basitrichous haplonemas and microbasic mastigophores can be detected with high DAPI concentrations (Sup. Fig. 4C,C’). An antibody against the minicollagen NvNcol-3, which recognises all three different cell types (Zenkert et al., 2011), was used to mark early developing nematocytes (Sup. Fig. 4D,D’). In parallel, we dissociated individual tentacles, and using DIC microscopy, we assessed all three types of nematocytes by morphology. All types were present in the *twist^−/−^* and WT tentacles, and there were no apparent changes in cnidocyte development in the mutant (Sup. Fig. 4E-G, E’-G’). This suggests that *twist* has no direct role in cnidocyte development. Lastly, we generated single cell transcriptomes from *twist^−/−^* tentacles and compared them with tentacle WT libraries. The analyses suggested that the mutants do not have any missing cell types. When comparing the relative abundance of each cell type, normalized to the total of cells per sample, we observed a decrease in spirocytes, muscle cells, neurons and mucus glands. Yet, the reduction in muscle cells may reflect the smaller tentacle size observed in *twist*^−/−^ mutants and could contribute to their altered morphology when compared to the WT (Sup. Fig5B).

The morphological differences between *twist*^−/−^ and WT animals become more pronounced as animals grow. In the *twist^−/−^* mutants, primary tentacles, much like initially secondary tentacles, get shorter and thinner, in addition, merged secondary tentacles are frequently seen, suggesting that proliferation deficiencies in the tentacles may continue throughout growth of the polyp. Additionally, in contrast to the essentially flat mouth ends of the WT, the initial light trapezoid-like structure seen in the primary polyp oral disc becomes more apparent in the adults. (Fig. 2G), (Fig. 6A,B,C; Suppl. Fig. 3B.

**Figure 6.**
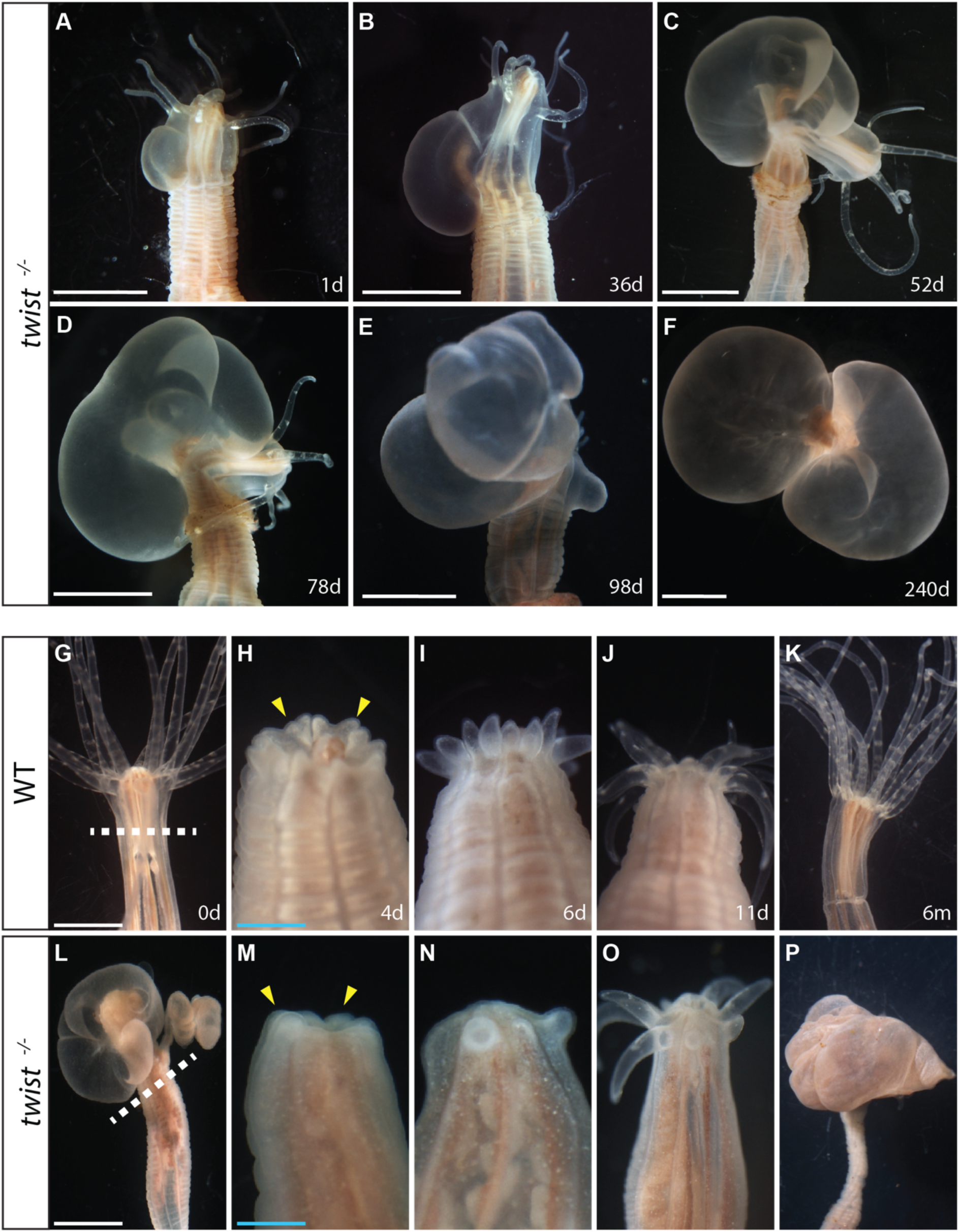
Impaired tentacle regeneration and neoplasm formation in *twist^−/−^* mutants. **(A)** Day 1 of neoplasm growth documentation **(B)** Day 36 **(C)** Day 52 **(D)** Day 78 **(E)** Day 98 **(F)** Day 120**. (G-K)** WT and **(L-P)** *twist^−/−^*animals at various time points following oral amputation (cut indicated by dashed line in G and L). **(H, M)** Four days post-amputation: yellow arrow heads mark the appearance of new tentacle buds. **(I, N)** Six days post-amputation: WT animals display synchronous regrowth of all tentacles, while *twist^−/−^* animals show limited and asynchronous tentacle regeneration. **(J, O)** Ten days post-amputation: WT animals continue regeneration tentacles uniformly; *twist^−/−^* animals exhibit uneven tentacle development. **(K, P)** Six months post-amputation: WT have regenerated all 16 tentacles, while *twist^−/−^* display a reappearance of a neoplasm. **(L)** Neoplasm is visible at the oral pole prior to amputation in twist^−/−^ animals. Scalebars: 5 mm white, 2.5 mm blue.

### Juvenile *twist^−/−^* mutants develop a neoplasm-like formation at the level of the pharynx

Most strikingly, in *twist*^−/−^ mutants starting at the juvenile stage (about 1.5 months postfertilization), a bubble-shaped neoplasm-like epithelial structure will begin to randomly appear on one side of the animal between two mesenteries. This expansion, which includes both cell layers, is mostly found in the *twist*-expressing region of the body wall at the level of the pharynx. The growth of the epithelial neoplasm starts at the level of the lower pharynx and later expands somewhat down the body column, away from the oral pole. It is dependent on feeding and will initially grow at a rapid pace slowing down in later stages. Since the growth of the neoplasm impacts the mouth region, the decrease in rate of tissue expansion could be an indirect result of a lower food intake. If left unchecked the neoplasm will eventually engulf the whole head of the animal. The animals can survive for some time in such a terminal state but will eventually die due to their inability to feed (Fig. 6E,F).

We next wished to know whether the two phenotypes – impaired secondary tentacle formation and epithelial neoplasm – are reproducible even after decapitation and regeneration. If the neoplasm is removed by bisection of the whole head, animals will regenerate a head and pharynx (Fig. 6L-O). Both WT animals and *twist^−/−^ mutant*s initiate tentacle regeneration simultaneously, yet, while WT animals regenerate all 16 tentacles at once, *twist^−/−^* mutants initially regenerate only 4 to 8 tentacles, reproducing the initial phenotype (Fig. 6 H,I,M,N). Later, the mutants may form an 2-4 additional, malformed or crippled tentacles (Sup. Fig. 3B & Fig. 6I). Thus, the program to generate tentacles in every other mesenterial segment is not affected in the mutant, but the formation of additional tentacles and the subdivision in pseudo-segments is also impaired during regeneration. Moreover, after about 1 month, also the epithelial neoplasm will reappear (Fig. 6P). These observations strongly suggest that the defects are reproducible and robust and, hence, likely to be of genetic cause.

To test whether the neoplasm is due to excessive proliferation or to a balloon-like inflation of the epithelial tissue, we conducted an EdU labelling experiment and compared the side of the animal with the neoplasm vs the normal side within the same mutant polyp. We examined 16 samples and found that the neoplasm had a significantly higher proliferation rate than the unaffected tissue (Fig. 7C). Sagittal sections of the neoplasm area revealed that the mesoglea, which separates the gastrodermis and ectoderm, becomes considerably thicker during body wall expansion. Notably, in addition to the gastrodermal circular muscle, the sections also revealed an ectopic ectodermal muscle. This is remarkable, as usually only tentacles have ectodermal muscles (Fig. 8A,B) (i.e. the longitudinal retractor muscles) (Jahnel et al., 2014). When investigated in more detail, we found that the ectopic muscle layer was longitudinally aligned, mirroring the orientation of the ectodermal muscle layer in the tentacles (Fig 8C,D).

**Figure 7.**
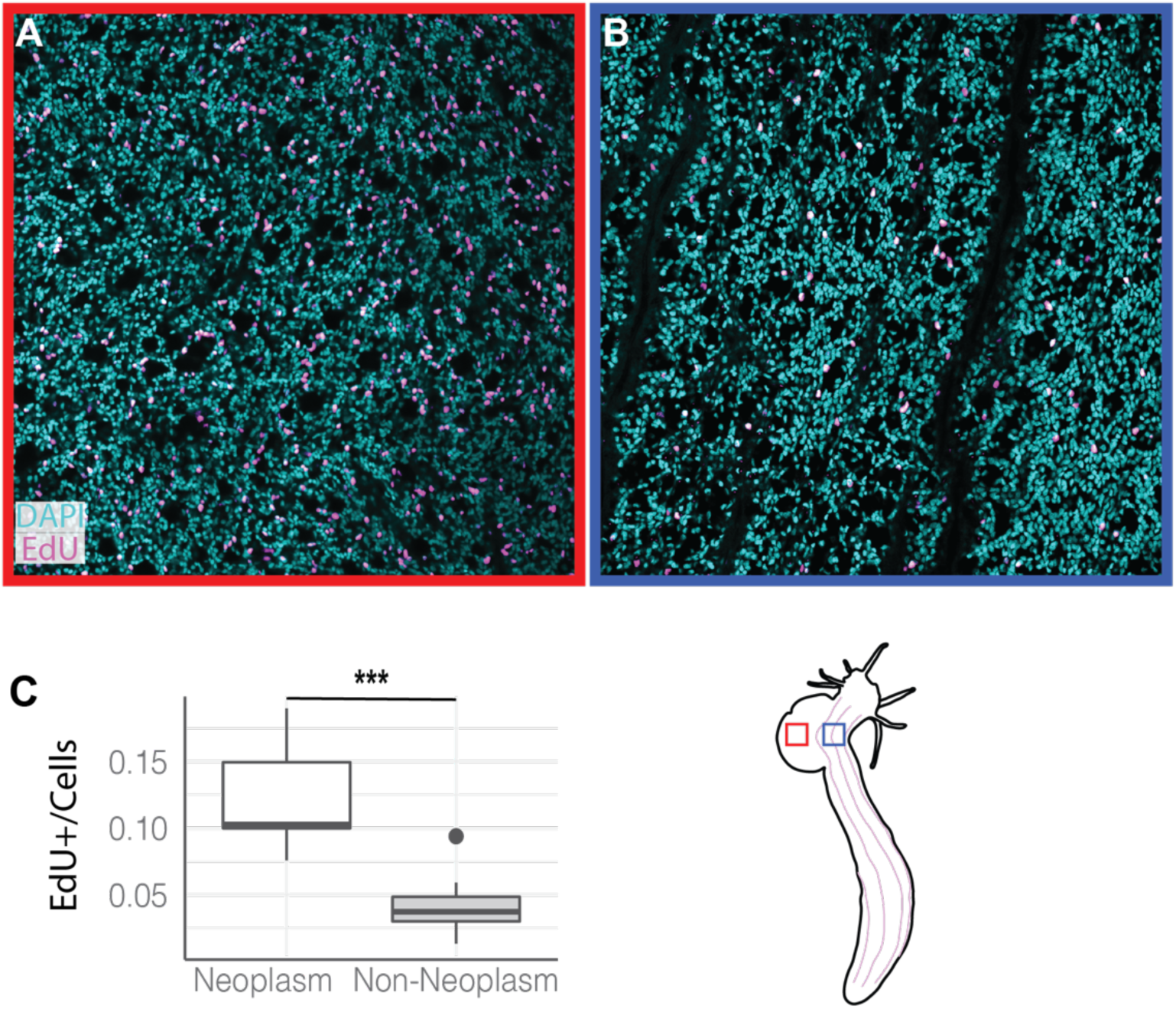
EdU Staining shows increased proliferation of neoplasm tissue in *twist^−/−^* animals. **(A)** EdU (magenta) and DAPI staining (cyan) of the neoplasm side, showing actively dividing cells in the expanding region. **(B)** EdU and DAPI staining of the non-neoplasm side, showing less cell division. **(C)** Quantification of the ratio of EdU-positive cells to total cell nuclei, indicating a higher cell division ratio on the bubble side compared to the non-bubble side. Scalebar is 50 µm. The accompanying sketch illustrates the sample locations: red for neoplasm and blue for non-neoplasm.

**Figure 8.**
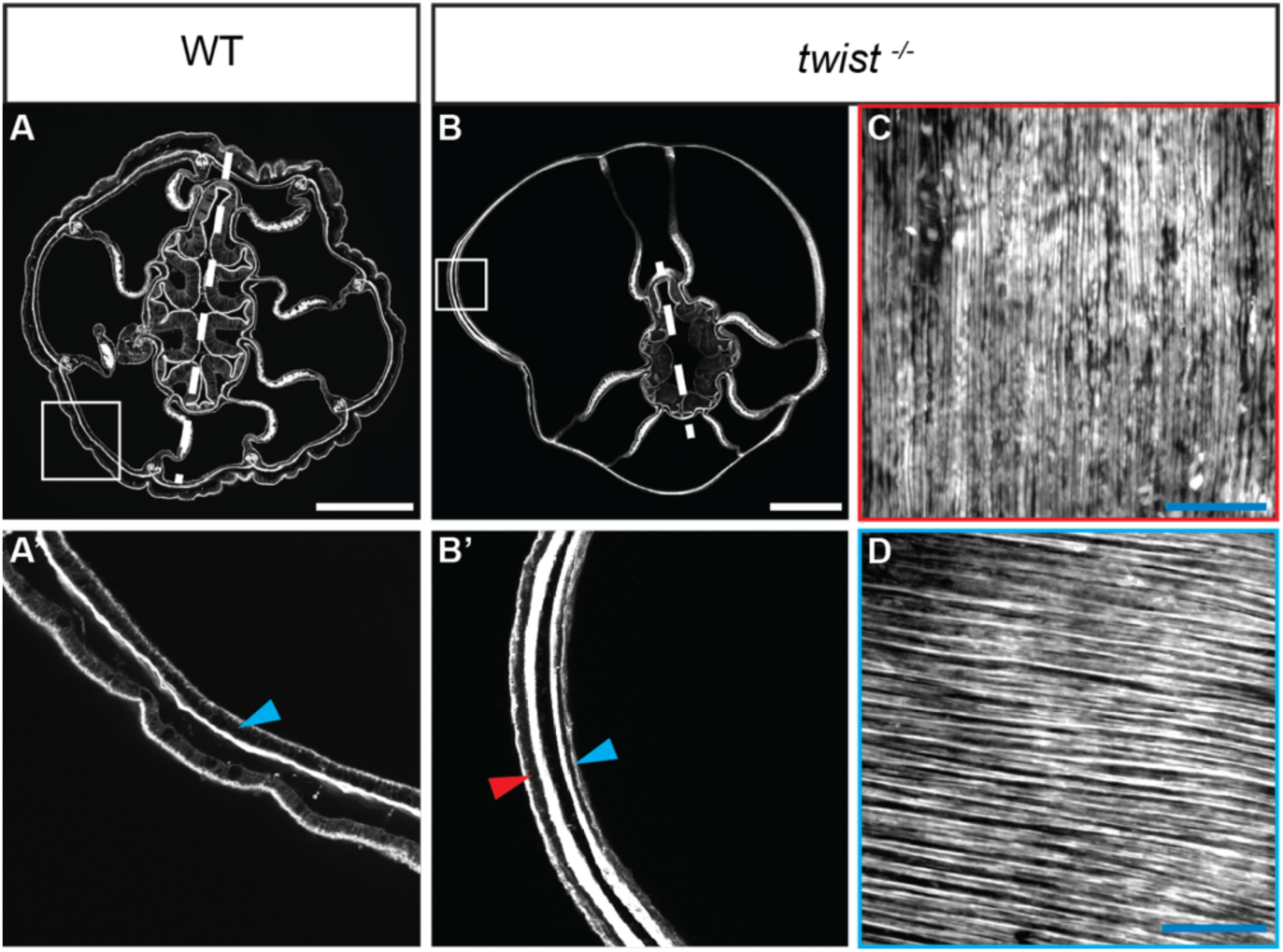
Sagittal sections of the neoplasm reveal ectopic longitudinal ectodermal muscle cells in *twist-/-* animals. **(A**) Phalloidin staining of a sagittal cross-section of an adult WT (wild type) at the height of the pharynx, showing the anatomical features of the region. **(B)** Phalloidin staining of a sagittal cross-section of an adult twist mutant at the height of the pharynx, highlighting the morphological differences compared to WT. **(A’)** Close-up view of the inlet shown in (A), with a blue arrow indicating the mesodermal muscle layer. **(B’)** Close-up view of the inlet shown in (B), with a blue arrow indicating the mesodermal muscle layer and a red arrow indicating the ectodermal muscle layer. **(C)** Horizontal close-up view of the ectodermal muscle layer of the neoplasm, revealing its longitudinal alignment. **(D)** Horizontal close-up view of the mesodermal muscle layer of the neoplasm, showing its horizontal alignment. Dashed lines are used to indicate the directive axis. The white scale bars represent a length of 500 µm, while the cyan scale bars represent a length of 50 µm.

This suggested that the neoplasm is possibly not only an epithelial overgrowth but may have adopted a tentacle-like identity. To test this, we generated scRNA-seq libraries from dissected neoplasm tissue and compared them to wildtype tentacle and body wall samples. We then clustered the combined dataset and annotated major cell types based on known markers (Fig. 9A). While the UMAP projection (Fig. 9A) does not distinguish cells by sample origin, analysis of the cell type distribution revealed that spirocytes (SP) and tentacle retractor muscle (TR) cell types were absent from the interparietal tissue of the wt, but present in the both the neoplasm and tentacle samples (Fig. 9B). Consistently, tentacle-specific transcripts such as *nem64* (a TF associated with the retractor muscle; (Cole et al., 2023), *avil-like1* (tentacle-specific ectoderm marker) and *ncol* (a spirocyte specific minicollagen) were expressed exclusively in the neoplasm and in tentacle samples, but not in the WT-interparietal control sample (Fig. 9C). We also observed an up regulation of *tbb-like1*specifically in the neoplasm-interparietal tissue (Fig 9C). While this beta-tubulin isoform is broadly expressed across nearly all clusters in the wildtype, it shows an enrichment in a subset of neural subtypes (Sup. Fig. 6) Together with alpha-Tubulin, beta-Tubulin forms the essential component of microtubules. Although the specific role of *tbb-like1* in the neoplasm is unclear, its upregulation may be related to the higher rate of cell division and the transformation into a neoplasm (see discussion). These data suggest that in juvenile *twist* mutants a neoplasm-like structure emerges by epithelial hyperproliferation, driving a differentiation of a tentacle-like profile.

**Figure 9.**
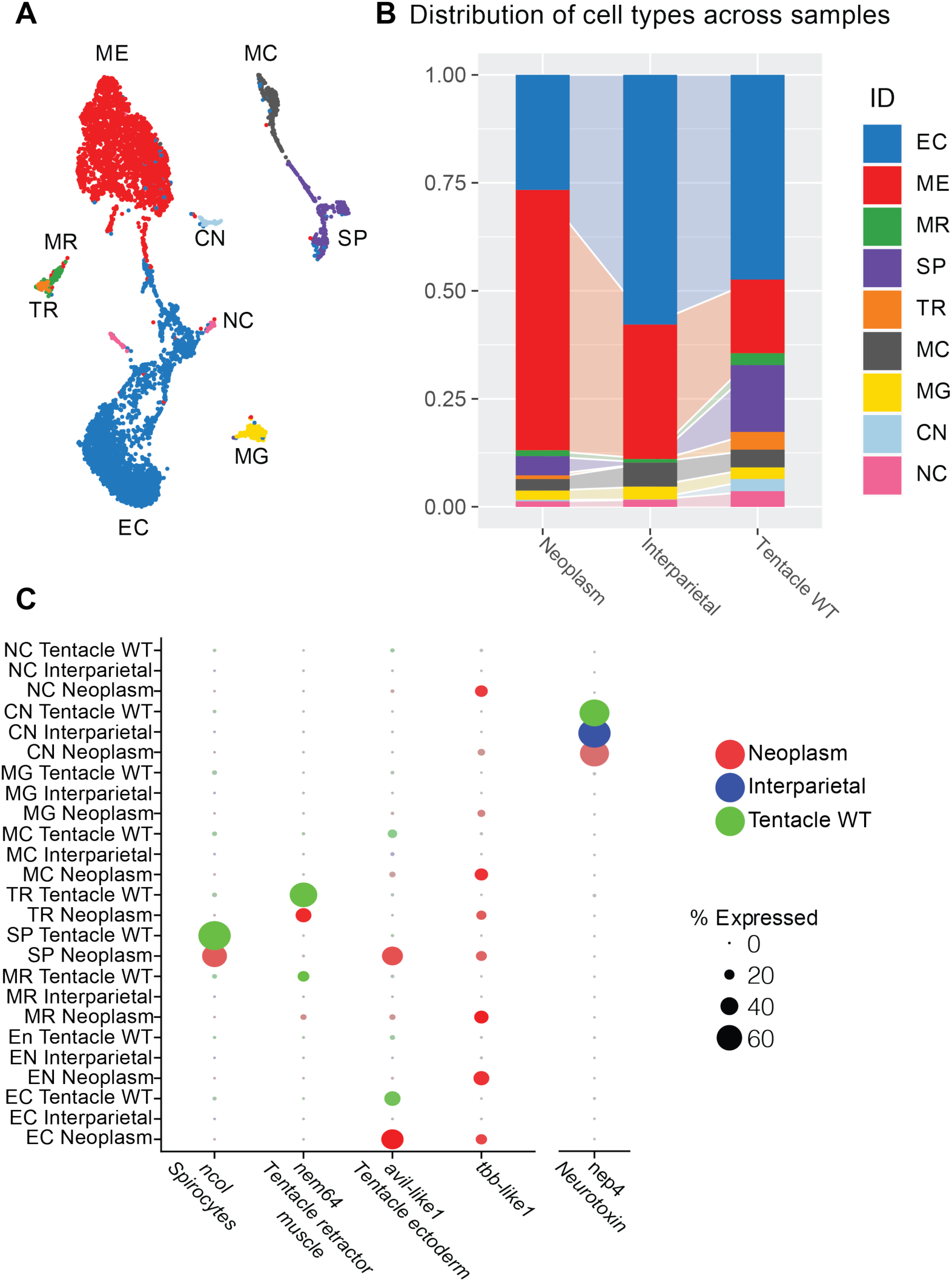
scRNA-seq analysis of neoplasm reveals a cellular profile reminiscent of tentacles. (**A**) UMAP projection of integrated single-cell transcriptomes from neoplasm, tentacle, and interparietal body wall tissues. Cell clusters were annotated based on known marker expression. Abbreviations: ME – mesoderm; EC – ectoderm; MG – mucus glands; CN – cnidocytes; MC – mature cnidocytes; SP – spirocytes; MR – mesentery retractor muscle; TR – tentacle retractor muscle; NC – not categorized. **(B)** Cell type composition of each tissue, showing that spirocyte (SP) and tentacle retractor (TR) clusters are absent in the interparietal sample but present in both the neoplasm and tentacle. **(C)** Dot plot showing expression of tentacle-specific genes (*ncol*, *nem64*, *avil-like1*) in neoplasm and tentacle samples. *tbb-like1* is specifically upregulated in the neoplasm, while *nep4* (a general cnidocyte marker) is expressed across all three tissues.

## Discussion

### *twist* in *Nematostella vectensis* is not essential for initial germ layer specification and mesentery segments

The Twist family of basic helix-loop-helix (bHLH) TFs was originally characterized in *Drosophila melanogaster* for mesoderm specification and muscle development (Leptin and Grunewald, 1990; Thisse et al., 1987; Thisse et al., 1988). It has been identified in a wide range of phyla from Cnidaria to vertebrates (Corsi et al., 2000; Martindale et al., 2004; O’Rourke and Tam, 2002; Spring et al., 2000; Wu et al., 2008; Yasui et al., 1998). Like all bHLH TFs, Twist has a highly conserved DNA-binding domain and two amphipathic helices essential for dimerization (Ellenberger et al., 1994) and a WR domain, which shows identical amino acid sequences between vertebrates and cnidarians but varies among lancelets, flies, and nematodes (Wu et al., 2008). Despite these highly conserved domains Twist displays a remarkable functional plasticity across different metazoan lineages (Davis et al., 1990; Ma et al., 1994). Central to this functional diversity is the ability of Twist to form homo-or heterodimers, as it is an obligate dimer and cannot bind to DNA as a monomer (Bouard et al., 2018; Chang et al., 2015; Maia et al., 2012). This dimerization capacity allows Twist to either activate or repress target genes according to cellular context, interacting partners, and stage of development (Castanon et al., 2001; O’Rourke and Tam, 2002). Consistent with this plasticity, our findings show that in *Nematostella vectensis*, *twist* expression is first detected at the post-gastrulation planula stage between 2 and 3 days of development (Martindale et al., 2004), within the developing mesodermal compartments surrounding the involuted pharynx, which later form the mesenteries. Unlike in *Drosophila,* knockdown of *twist* does not impair gastrulation, thus, does not seem to have a crucial role in germ layer specification, indicating that Twist has no essential role in this process. Additionally, Twist is also not required for the formation of the first eight mesentery segments, which are under genetic control of the BMP signalling gradient and the downstream Hox genes (Saina et al., 2009; Genikhovich et al., 2015; He et al., 2018).

While this pattern is distinct from *Drosophila*, it is consistent with observations in other organisms, such as vertebrates, nematodes, and lancelets, in which *twist* is expressed after gastrulation and is not required for germ layer specification (Harfe et al., 1998; Molkentin et al., 1995; Spring et al., 2000; Yasui et al., 1998). Consistent with this, *twist* is not among the early target genes of signaling pathways specifying the three germ layer identities in *Nematostella* (Haillot et al., in press). Interestingly, in another winged insect, *Tribolium castaneum, twist* is required for mesoderm formation but unlike in *Drosophila* it does not function as the sole determinant, as mesoderm continues to be specified throughout segmentation (Handel et al., 2005). This suggests that the strict dependence on Twist for mesoderm specification in *Drosophila* is a derived feature, rather than a general trait of insects or bilaterians. Other studies have highlighted a role of Twist in epithelial-to-mesenchymal transition (EMT) during gastrulation, neural crest delamination or metastasis (Fan et al., 2021; Qin et al., 2012; Yang et al., 2004; Chen and Behringer, 1995; Thisse et al., 1987). While gastrodermal (mesodermal) cells in *Nematostella* undergo a partial EMT during early gastrulation (Kraus and Technau, 2006; Pukhlyakova et al., 2019), our data do not support an ancestral role of *twist* in EMT. These findings continue to highlight the diverse and species-specific functions of Twist across metazoans.

### Twist regulation by Wnt, Notch and BMP signalling in *Nematostella vectensis*

In vertebrates, Wnt signalling plays a major role in neural crest development, where it regulates multiple TFs, including Twist (Ji et al., 2019). Additionally, Wnt-induced *twist* expression has also been observed in other contexts, such as cranial mesenchyme during bone formation, in chondrogenesis, where it functions to inhibit cartilage differentiation, and tumour progression (Howe et al., 2003; Reinhold et al., 2006; Goodnough et al., 2016; Sun and Liu, 2017; Ji et al., 2019). While direct transcriptional regulation of *twist* by Wnt has been demonstrated in these contexts, evidence for *twist* activation by Wnt signalling outside of vertebrates remains scarce. However, our findings in *Nematostella* show that Wnt signalling is both necessary and sufficient for *twist* expression, suggesting an ancient role for this regulatory interaction. Whether the underlying molecular mechanisms are conserved across cnidarians and bilaterians remains an open question.

In addition to Wnt, both Notch and BMP signalling regulate *twist* expression in *Nematostella*. Our findings show that inhibition of either pathway results in a complete or near complete loss of *twist* expression. In *Drosophila*, Notch functions primarily as a modulator of *twist*, refining its expression during mesodermal patterning rather than acting as a direct activator (Tapanes-Castillo and Baylies, 2004). In contrast, in vertebrates, direct regulation of *twist* by Notch has been demonstrated in both development (Tian et al., 2015) and cancer progression (Wang et al., 2020). In *Nematostella*, our results indicate that Notch signalling is necessary for *twist* expression, yet it remains unclear whether this regulation occurs through direct activation, as seen in vertebrates, or through modulation, as observed in *Drosophila*. Wnt and Notch signaling form a feedback loop during early germ layer specification in *Nematostella* (Haillot et al., in press), therefore, the loss of *twist* expression upon inhibition of Notch signaling may be due to its role in regulating Wnt signaling.

BMP signalling has a well-established role in axial patterning and mesentery formation in *Nematostella* (Saina et al., 2009; Genikhovich et al., 2015), and our findings indicate that BMP signalling is necessary for *twist* expression. While the underlying molecular mechanism remains unclear, studies in vertebrates suggest that BMP can modulate *twist* expression in a context-dependent manner. In chick embryos, BMP2 has been shown to downregulate or upregulate *twist* expression in the paraxial mesoderm depending on the developmental stage (Hornik et al., 2004). Whether BMP modulates *twist* expression in a similar developmental-stage-dependent manner remains to be determined, and further work is needed to assess whether BMP-*twist* interactions reflect conserved regulatory logic or lineage-specific mechanisms.

In contrast to Wnt, Notch and BMP, inhibition of MAPK had no effect on *twist* expression, indicating that this pathway does not play a major role in *twist* activation in *Nematostella*. However, studies in vertebrates have shown MAPK can act upstream of Twist through post-translational mechanisms. For example, in mammalian systems, Twist1 is phosphorylated by MAPK pathways, including ERK, JNK, p38 and MEK1/2 preventing its ubiquitination and subsequent degradation (Hong et al., 2011; Li et al., 2013). This leads to increased Twist1 protein levels, which enhances its function during EMT and tumour invasion. While such mechanisms have not been described in *Nematostella*, it is worth noting that in the annelid *Platynereis dumerilii* regions of high MAPK activity overlap with *twist* expression during early development, hinting at a possible regulatory interaction, although a direct link was not shown (Pfeifer et al., 2014). Together, these findings suggest that while MAPK does not regulate *twist* transcription in *Nematostella*, it may influence Twist activity through post-translational or context-dependent mechanisms in other lineages. Whether such regulation also occurs in early-branching animals remains to be investigated.

Inhibiting Hedgehog (SHH) had no effect on *twist* expression in *Nematostella*, indicating that *twist* is not downstream of SHH signalling. However, in vertebrates the regulatory relationship between Twist and SHH is more complex and context dependent. In mouse, loss of Twist1 leads to reduced *Shh* expression in the limb bud, likely due to failure of apical ectodermal ridge (AER) formation and subsequent loss of FGF signalling (O’Rourke and Tam, 2002). At later stages Twist1 acts a repressor of *Shh* expression, preventing ectopic activation during limb development (Zhang et al., 2010). In chick embryos, *twist* expression requires the combined input of FGF8 and SHH, supporting a role of Twist as an integrator of multiple upstream signals (Hornik et al., 2004). In addition, Twist1 influences the expression of downstream SHH pathway components such as *Gli1*, *Gli2*, *Ptch*, and *Hand2*, suggesting a broader regulatory role within the SHH network (Qin et al., 2012). Recent evidence suggests that Twist can also regulate SHH pathway output by activating *Gli1* transcription in specific contexts (Tu et al., 2024).

In *Nematostella, paraxis* and *tbx15* expression overlaps with *twist* and both genes are strongly downregulated in the *twist^−/−^* mutants, indicating that Twist is required for their normal expression. *Paraxis* belongs to the Twist-family of bHLH TFs, and its expression overlaps with *twist* and often plays a similar developmental role in bilaterians, including vertebrates and spiralians (Wilson-Rawls et al., 2004; Barnes and Firulli, 2009). In the brachiopod *Terebratalia*, *paraxis* is expressed after *twist*, suggesting a possible conserved regulatory relationship in which Twist functions upstream of *paraxis* (Passamaneck et al., 2015). In contrast, the relationship between Twist and T-box genes remains less straightforward. For example, in chick, *twist*, *tbx4/5* and *snail* are co-expressed in the limb and are induced by FGF signalling, yet appear to function in a parallel rather than in a hierarchical manner (Isaac et al., 2000). In contrast to *Nematostella*, in *Drosophila*, *twist* is downstream of the Tbx1 ortholog *Org-1* (Rose et al., 2022), raising the possibility of a lineage specific regulatory relationship or a regulatory interaction that has diverged over time.

### Twist controls the morphogenesis of secondary tentacles

A significant finding was that juvenile *twist* mutants had abnormal oral disc morphology and shorter tentacles. Previous research established stereotypical phases of tentacle addition in *Nematostella* (Fritz et al., 2013; Ikmi et al., 2020). Twist was discovered to be necessary for the timely emergence and morphogenesis of secondary tentacles during post-embryonic development, while primary tentacle formation remains largely unaffected (besides minor differences in size after longer development). This suggests that the primary tentacles are still governed by an embryonic/larval program, which does not involve Twist function, while secondary tentacle formation requires new patterning and induction. The *twist* mutation resulted in delayed and aberrant tentacle formation, a phenotype reminiscent of Hydra regeneration-deficient mutants in which impaired Wnt/β-catenin signalling correlates with failure of head and tentacle development (Hobmayer et al., 2000). The shorter tentacle phenotype may also be a result of continued diminished proliferation. Our findings suggest that Twist is essential for the proper signalling dynamics between TOR and FGF pathways, which are critical for normal tentacle growth (Ikmi et al., 2020). In *twist* mutants, we observe delayed activation of pRPS6 spots in the early tentacle buds indicating impaired TOR signaling activation. Additionally, while *Fgfrb* expression was initially detectable in both WT and *twist* mutants, its expression weakened as *twist* mutants matured, particularly at the initiation of secondary tentacle budding. Given that *twist* is expressed in the gastrodermal layer and pRPS6 signaling is initially active in the ectodermal layer, the activation of the TOR parthway is likely mediated indirectly through FGF signaling. This aligns with the role of FGFRb in local activation the TOR pathway, as demonstrated before (Ikmi et al., 2020).

Notably, *twist* mutants fail to form micronemes, which are incomplete mesentery folds from the body wall marking the middle of each segment. This subdivision of the segments predicts the sites of new tentacle appendages in the wildtype. The lack of the micronemes indicates a disruption in directive axis patterning (Ikmi et al., 2020). It has been previously shown that proper formation of the eight mesenterial segments is crucial for position and development of the primary tentacles in even-numbered segments (He et al., 2018). Given this context, it is feasible that the lack of micronemes in the *twist* mutant is associated to the observed reduction in the number of secondary tentacles. Micronemes, which delineate boundaries between neighbouring tentacles within the same segment, likely play a critical role in the spatial organization necessary for proper tentacle development. The absence of these structures in the *twist* mutant could lead to impaired segmentation, resulting in fewer and improperly positioned secondary tentacles. Taken together, this suggests that Twist operates at the tissue/morphogenetic level but is not indispensable for all differentiation programs. However, the casual connection between the lack of micronemes and the impaired regulation of localized TOR and FGF signaling remains unclear at this point. Future research should aim to elucidate the transcriptional targets and signals that Twist regulates during tentacle development.

### Distinct Roles of Twist in Muscle Development Across Species

Our findings indicate that Twist in *N. vectensis* does not impact muscle development or specification, despite the morphological abnormalities, such as shortening, thinning, and fusion observed in the *twist* mutant tentacles. The appropriate formation and orientation of muscle layers within these altered structures contrast starkly with functions of Twist in other organisms. For instance, the nematode *twist* ortholog, *hlh-8*, is integral to the differentiation of certain mesoderm-derived muscle lineages (Corsi et al., 2000). Similarly, Twist functions in mice as a repressor, staving off premature differentiation of myogenic cells and ectopic myogenesis (Bialek et al., 2004). In the hydrozoan *Podocoryne carnea*, *twist* was proposed to have a potential role in striated muscle formation of the medusa, because it is expressed in the entocodon, which contributes to the differentiation of smooth and striated muscle cells (Spring et al., 2000). However, in the related hydrozoan *Clytia hemisphaerica, twist* was shown to be expressed in stripes adjacent to the radial canals, non-overlapping with the striated muscles (Kraus et al., 2015). This suggests that like in *N. vectensis*, *twist* does not appear to be essential for muscle formation. This view is reinforced by recent research demonstrating that the bHLH transcription factor, Nem64, is necessary for formation of the ectodermal tentacle retractor muscle in *N. vectensis* (Cole et al., 2023). This discovery highlights the species-specific roles of Twist and its related factors, shedding light on the intricate process of muscle cell type diversification caused by the duplication of bHLH transcription factor genes.

### *twist* mutant induces oral body wall neoplasms have a tentacle-like profile

Interestingly, elder juvenile *twist* mutants developed oral body wall neoplastic outgrowths. These epithelial neoplasms displayed increased proliferation and surprisingly single cell RNAseq showed a cellular and molecular profile reminiscent of tentacles. This included the presence of spirocytes, which is a tentacle-specific cnidocyte cell type and of the tentacle-specific retractor muscle gene *nem64*. In line with that, we detected longitudinal musculature characteristic of the ectodermal tentacle lineage. This is quite surprising, as *twist* mutants first fail to locally activate proliferation thereby inhibiting secondary tentacle formation. This is unexpected, as *twist* mutants initially fail to locally activate proliferation, thereby inhibiting secondary tentacle formation. The emergence of unpatterned, proliferative tissue with tentacle-like differentiation suggests that Twist plays a crucial role in spatial control during post-embryonic development not only promoting tentacle formation in appropriate domains but also suppressing it in others. These findings suggest that while the neoplasm adopts a transcriptional tentacle-like identity, it fails to undergo proper morphogenesis. This points to a possible uncoupling of tissue identity from shape, where the appropriate cell types are specified but their spatial organization is lost. In this context, the micronemes, which are absent in *twist* mutants, may function not only as signalling hubs but also as physical landmarks that constrain and guide tentacle morphogenesis. Their loss could remove both molecular cues and spatial boundaries necessary for localized outgrowth, leading to the disorganized tentacle-like structures observed in the mutant. One marker gene upregulated specifically only in the neoplasm but not expressed in the tentacles nor the wildtype body wall is *tbb-like 1*, a beta-tubulin gene. This may reflect an increased rate of cell proliferation; however, its high specificity suggests a more defined role. Notably, beta-tubulin isoforms are upregulated in various cancer types, with some studies reporting **nuclear localization** and **functions** independent of microtubules (Walss-Bass et al., 2002; Ruksha et al., 2019; Kanakkanthara and Miller, 2021; Ludueña et al., 2022). Although their precise function in tumorigenesis remains unclear, tubulin isoforms are considered promising drug targets, hinting at an important role in neoplastic transformation. Future work should investigate whether *tbb-like1* has a similar tumorigenic role in the formation or maintenance of neoplasms in *twist* mutant *Nematostella*. Notably, tumors have rarely been reported in cnidarians. One example is a naturally occurring germline tumor in *Hydra* (Domazet-Loso et al., 2014), which is transplantable and independent of the cellular environment. A transcriptomic analysis of these tumours identified 44 tumour-specific genes; comparison with our neoplasm data revealed no overlap with *tbb-like1*. However, the gene *TCTP-like-1*, a known cancer-associated factor, was upregulated in both datasets (Domazet-Lošo et al., 2014; Sup. Table 1). The *Hydra* tumor is thought to result from defective differentiation of gamete-restricted stem cells, while the neoplasm observed in *twist* mutants appears to originate from epithelial cells. Interestingly, similar neoplasm-like outgrowths were recently reported in *Nematostella* with CRISPR/Cas9-induced *bmp2/4* knockout, where bubble-like structures formed near the oral pole (Knabl et al., 2024). Determining the transcriptional targets and signalling interactions regulated by Twist particularly with pathways like BMP will be essential to uncover how body patterning and homeostasis are maintained, and how their breakdown leads to neoplasia.

## Methods

### Animal husbandry

The *Nematostella* polyps were maintained under the conditions outlined in previous studies (Fritzenwanker and Technau, 2002; Genikhovich and Technau, 2009b) at a temperature of 18°C and predominantly in the absence of light. The induction of spawning was facilitated through the combined influence of light and increased temperature, (Fritzenwanker and Technau; 2002). The embryos were incubated at a temperature of 21°C.

### CRISPR/Cas9 mutagenesis

CRISPR/Cas9 induced deletion was done as described (Ikmi et al., 2014). Using CRISPOR (Concordet and Haeussler, 2018) a guide RNA (gRNA) targeting the HLH domain of *Nematostella twist* (**CCC**ACGTTACCTTCCGATAAACT) was designed, synthesized with the MEGAshortscript T7 kit, and purified with 3 M sodium acetate/ethanol. 500 ng/µl of gRNA was co-injected into *Nematostella* embryos along with 1µg/l recombinant Cas9 protein (PacBio) and raised at 21°C. To determine the efficiency of the CRISPR/Cas9-induced deletion tentacle tips from injected primary polyps were taken and genomic DNA extracted (gDNA). Each tentacle tip was treated in 20 μl DNA extraction buffer (10 mM Tris-HCl pH 8, 50 mM KCl, 0.3% Tween 20, 0.3% NP40, 1 mM EDTA) with 1 μg/μl proteinase K (Thermo Fisher) at 58 °C, and inactivated at 98 °C for 20 minutes. A 2µl gDNA sample was used for melting curve PCR analysis. After the efficiency of the gRNA was confirmed, F0 injected animals were raised to sexual maturity and crossed with WT animals. F1 progeny with detected indels were sequenced and pooled by mutation type and interbred at sexual maturity to yield F2 homozygous mutants.

### Whole mount *in situ* hybridisation (WMISH)

WMISH carried out as described earlier (Genikhovich and Technau, 2009b), but with minor adjustments to improve the detection of the *twist* gene. Embryos were fixed in 4% PFA in Phosphate-Buffered Saline (PBS) for 1 hour at room temperature (RT), then washed 5 times with 0.2% Tween in PBS at room temperature. The embryos were then washed once with 50% methanol in PBS containing 0.2% Tween before being transferred to 100% methanol and kept at −20 °C until further processing. 1 ng riboprobe/µl hybridization was done overnight at 63°C, followed by detection with an alkaline phosphatase conjugated anti-digoxigenin (1:2000) antibody (Anti-Digoxigenin-AP, Fab fragments, Roche, REF 11093274910). Upon staining, the embryos underwent a series of five washes in PTw. Subsequently, the embryos were immersed in PBS containing 2% w/v triton-x100 until the background reached an acceptable threshold. The process was repeated until a robust signal was detected.

### Drug treatments

Embryos were incubated for 24 h in *Nematostella* medium containing the respective drug dissolved in DMSO. Treatments were predominantly done between 48 to 72 hours post fertilization (hpf), except for one batch treated with Azakenpaullone (AZ, GSK3β inhibitor, 5 µM; Sigma) from 24 to 48 hpf to assess potential early effects of Wnt signalling on *twist* expression. The following concentrations were used: ICRT14 (Wnt/β-catenin inhibitor, 50 µM; Sigma), Cyclopamine (Smoothened inhibitor, 10 µM; Sigma), U0126 (MEK inhibitor, 20 µM; Sigma), K02288 (BMP receptor inhibitor, 6 µM; Sigma), LY-411575 (Notch signalling inhibitor, 50 µM; MedChemExpress #HY-50752), and AZ. Control embryos were incubated in equivalent concentrations of DMSO under identical conditions. After incubation, embryos were fixed and processed for in situ hybridisation analysis as described above.

### Immunohistochemistry and staining

For EdU labeling, the Click-iTTM EdU Cell Proliferation kit (Invitrogen) was used. Primary polyps at the 5+6 tentacle bud stage were incubated for 30 minutes at RT with 50 M EdU in 16% ASW. The “bubble” and unaffected adult tissues, on the other hand, were treated for 1 hour at RT. All specimens were then anesthetized with 7% MgCl2 in 16% ASW. The primary polyps were fixed in 4% PFA in PBS for one hour at room temperature, while adult tissues were fixed overnight at 4°C. The tissues were then premetallized, blocked, and the Click-it reaction was performed following standard protocols. In cases where additional antibody staining was required, another round of permeabilization and blocking was done before an overnight incubation at 4°C with rabbit-anti-pRPS6 (Cell Signaling #4858). The tissues were then washed and counterstained overnight at 4°C with anti-rabbit secondary antibodies (1:2000 dilution) and 14 M DAPI in a buffer of PBS supplemented with 10% goat serum, 1% DMSO, and 0.1% Triton X-100. For phalloidin staining, polyps and adult tissues were fixed according to their developmental stage, then stained with 2.2M Alexa Fluor® 488 phalloidin for 2 hours at room temperature followed by five washes in PBT (0.5% Triton X-100 in PBS). Adult polyps were embedded in 10% gelatin in PBT and sectioned at 70-100 µm thickness with a Leica VT1200 vibrating blade microtome.

### Cnidocyte Staining

Primary polyps were anaesthetized with 7% MgCl2/16 ASW for 10–20 minutes, as per the protocol modified from (Wolenski et al., 2013). 4% FA/16% ASW/0.5M EDTA was used for a quick pre-fixation, and the same solution was used for a 1-hour fix at 4°C after that. Samples were rinsed with Tris-EDTA washing buffer (10 mM Tris, 10 mM EDTA, 10 mM NaCl, pH 7.6) at least three times after fixation. After that, they were incubated with 140 M DAPI/0.66 M Alexa Fluor® 633 Phalloidin in the washing buffer for a whole night at 4 °C. Prior to mounting, a series of five Tris-EDTA buffer washes were performed after incubation.

All samples were finally incubated overnight at 4°C in Vectashield® antifade mounting solution for maximum preservation and image quality.

### Single-cell RNA sequencing

Adult *Nematostella* tissues were prepared for single-cell RNA-seq, including tentacles, interparietal, and bubble (neoplasm). The tissues were treated with TrypLE^TM^ Select (Thermo Fisher, A1217701) for 4 hours to assist dissociation into a single-cell solution. The samples were gently pipetted up and down every 30 minutes to aid in dissociation, with extra care taken to minimize suctioning, especially during the first hour of the incubation. This procedure was repeated until a homogeneous single-cell suspension was obtained. Cell viability and counts were then determined using a Nexcelom Cellometer X2, and suspensions were diluted to 1000-2000 cells/µl. To ensure cell viability, the samples were stored on ice until they were ready for processing. The single-cell suspensions were put into a 10x Genomics single-cell platform, and libraries were produced using Vs.2 reagents according to the manufacturer’s instructions. The sequenced libraries were run through the CellRanger 5.1.0 pipeline with the default settings, with the goal of recovering 7000 cells per library. The reads were mapped and aligned to the most recent *Nematostella* genome (Zimmermann et al., 2023).

### Single-cell analysis

Single-cell transcriptomic analysis was performed using the Seurat R package v7.1 (Butler et al., 2018; Stuart et al., 2019). Raw UMI count matrices from libraries (“tentacle twist”, interparietal”, “neoplasm”, “tentacle WT”) were processed. Quality control filters retained cells with 200-2500 detected genes and fewer than 10,000 transcripts per cell. Highly variable genes were identified using the ‘vst’ method, and data were normalised with *LogNormalize*. Gene IDs were updated to *Nematostella vectensis* gene model annotations using a custom mapping table. To integrate datasets (“Tentacle twist”, Interparietal”, “Neoplasm”, “tentacle WT”), *FindIntegrationAnchors* and *IntegrateData* were used followed by scaling, PCA (50 components). Uniform manifold approximation and projection (UMAP) was used for dimensionality reduction. Clustering was tested over resolutions ranging from 0.1 to 0.8 with final analyses performed at 0.8, using shared nearest neighbour graph. UMAP *DimPlots* were used for library comparisons, *DotPlots* were used to illustrate expression patterns, and cluster identities were assigned based on marker gene expression. Differential gene expression between libraries was assessed using Seurat’s *FindMarkers* function (log2 fold-change > 0.25, detection rate > 25%).

## Acknowledgements

We would like to thank our animal facility staff, Angela Caballero, Vendula Stejskalova, Wolfgang Göschl, for continuous excellent care and support. This work was funded by a grant from the Austrian Science Fund FWF to UT (P34404). For the purpose of Open Access, the authors have applied a CC BY public copyright license to any Author Accepted Manuscript (AAM) version arising from this submission.

## Supplementary Figures

**Supplementary Figure 1.**
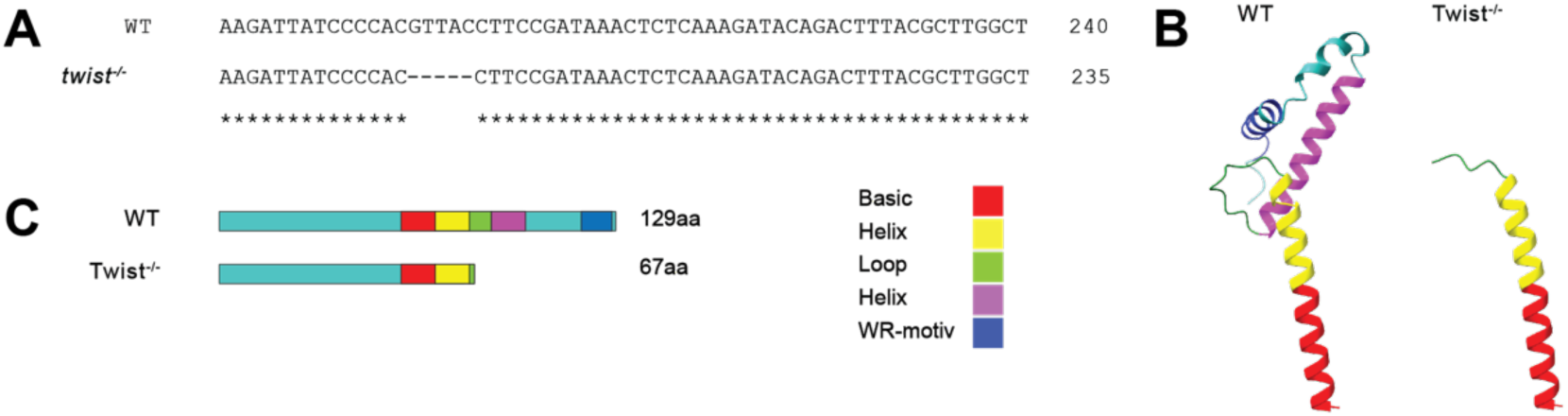
Sequence and structural comparison of WT and Twist^−/−^ proteins. **(A)** Segment alignment of WT and *twist^−/−^*nucleotide sequences showing a 5 bp deletion. **(B)** Alphafold predicted 3D structures of the bHLH domain of the WT and Twist^−/−^ proteins (Jumper et al., 2021). **(C)** Schematic representation of the WT and Twist^−/−^ proteins with colour-coded functional domains. WT protein is 129 amino acids long; Twist^−/−^ protein is 67 amino acids long.

**Supplementary Figure 2.**
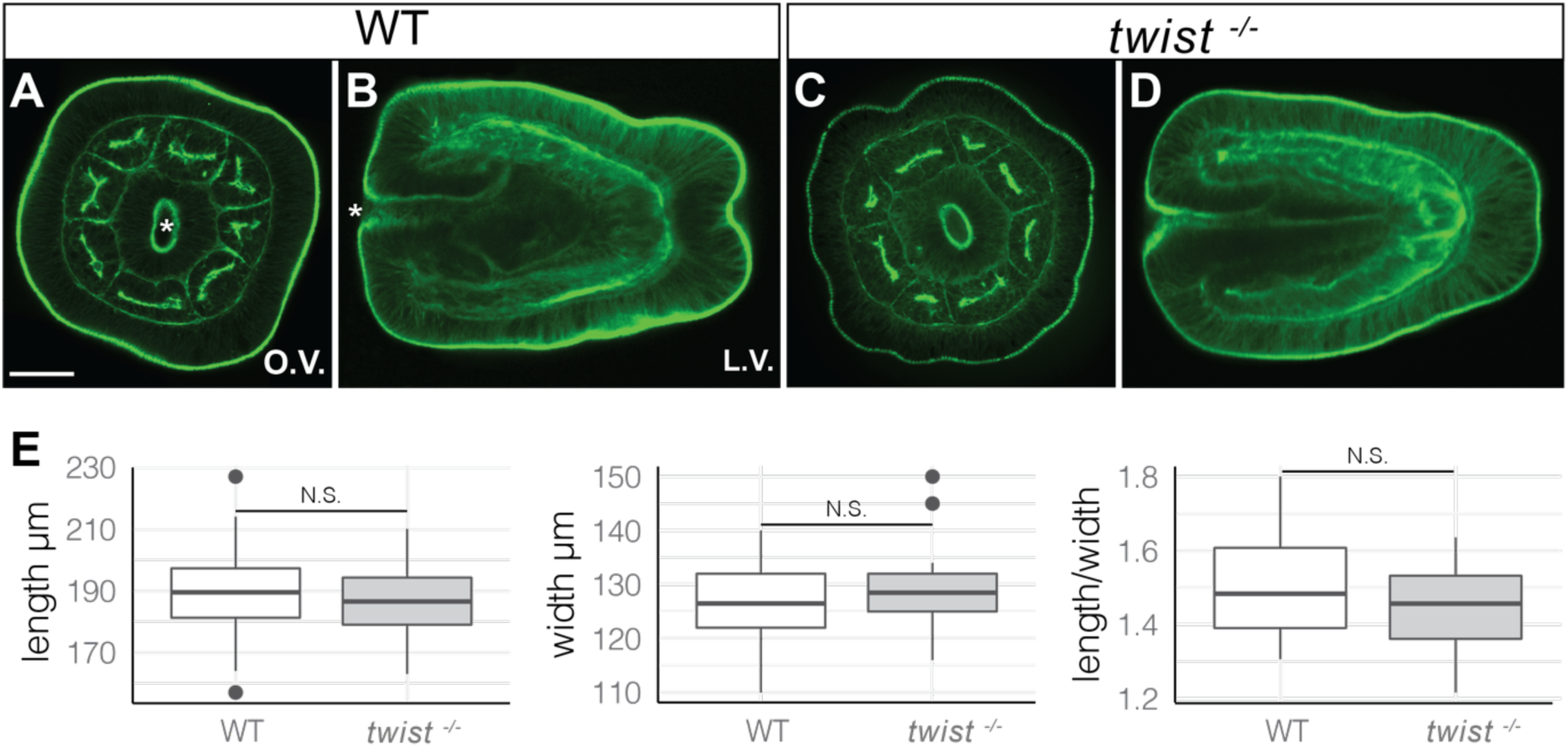
Pre-mesenterial segment development and quantification of body proportions in WT and *twist^−/−^*animals. **(A, B)** Oral and lateral confocal projections of a 4-day-old larva stained with phalloidin, showing the eight pre-mesenterial segments. **(C, D)** Oral and lateral views of a *twist^−/−^*larvae at the same stage. Asterisks mark the oral pole. **(E)** Box plots quantifying length, width, and length-to-width ratio for WT and *twist^−/−^* larvae. Scalebar: 50 µm.

**Supplementary Figure 3.**
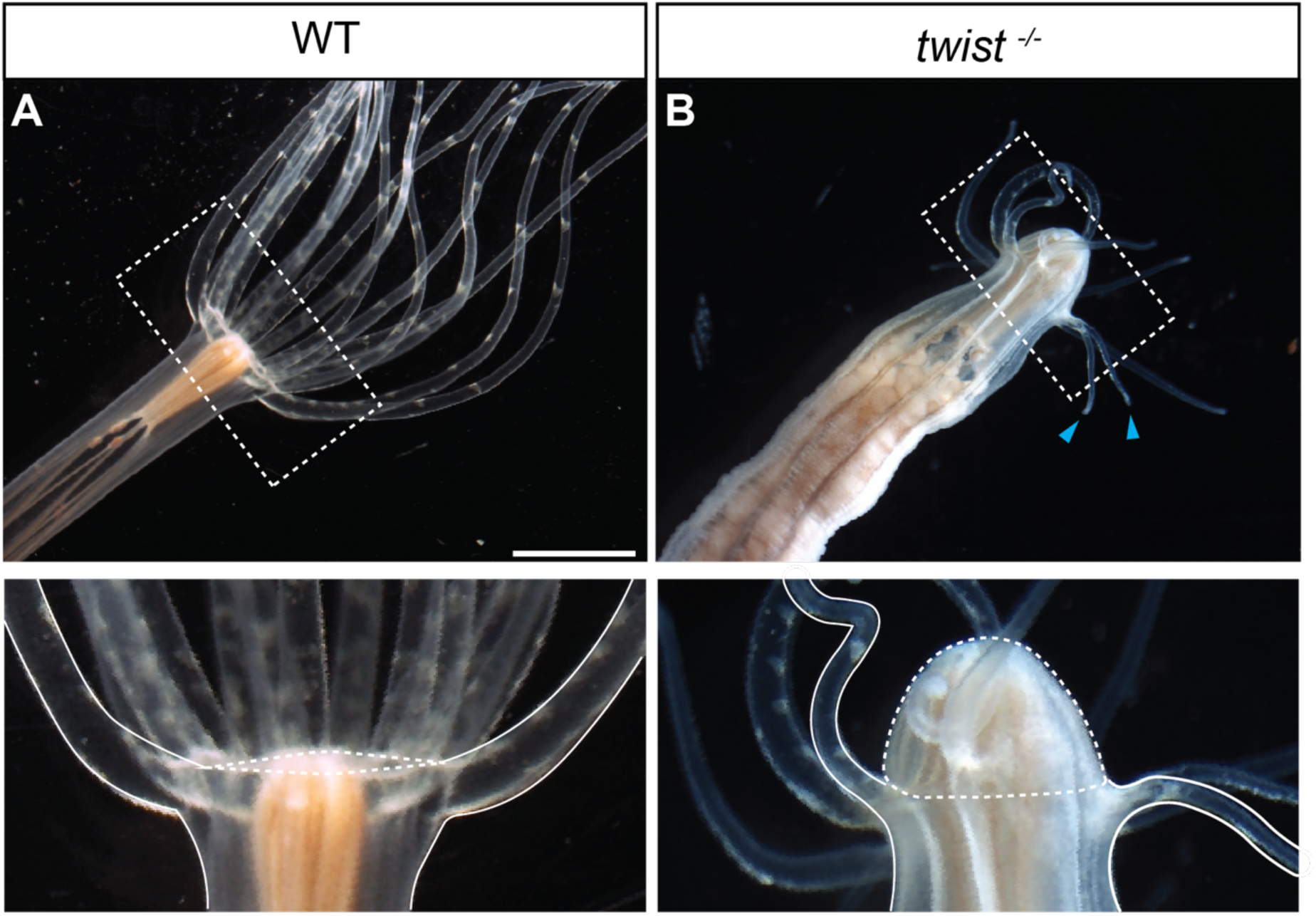
Adult tentacle number in WT and *twist^−/−^* animals. **(A, B)** WT polyps at 4 months (A) and 1 year (B), displaying typical formation of 16 tentacles. **(C, D)** *twist^−/−^*animals at the same developmental stages with reduced tentacle numbers, showing 5 (C) and 10 (D) tentacles respectively. Scalebars A, 2 mm; B, 5 mm.

**Supplementary Figure 4.**
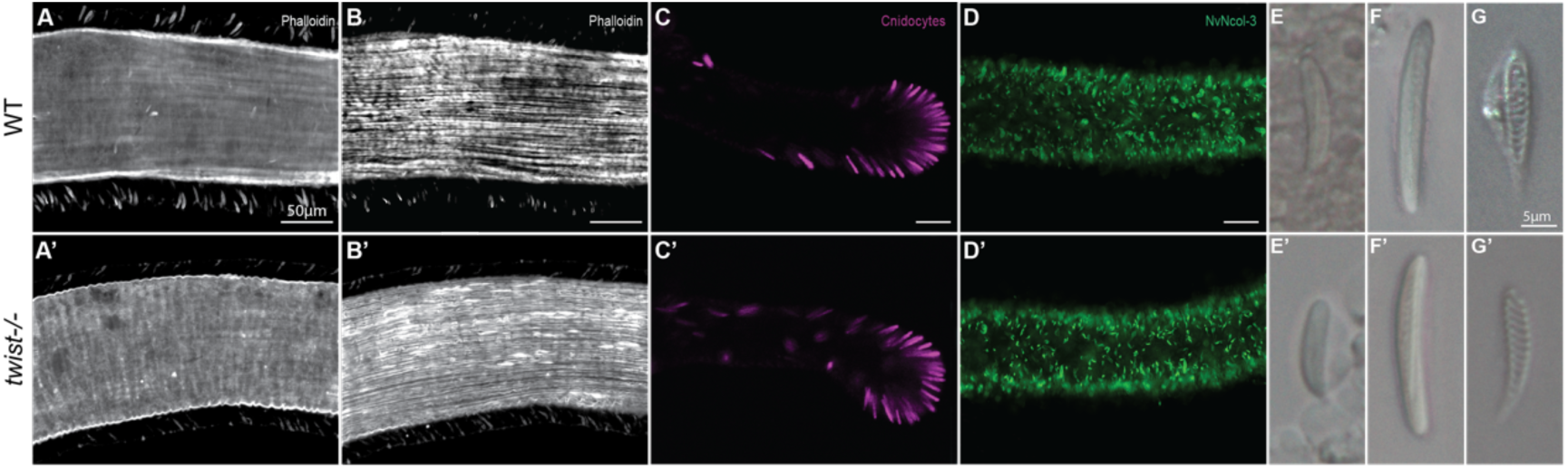
Muscles and nematocytes of WT and *twist^−/−^* tentacles. **(A, B)** Phalloidin staining of circular mesodermal (A) and ectodermal (B) tentacle muscles in WT animals. Ectodermal (longitudinal) and mesodermal muscles (circular) in *twist*^−/−^ mutants maintain their respective orientations **(A’, B’)**. **(C, C’)** DAPI staining (magenta) of Basitrichous Haplonemas and Microbasic Mastigophores present in both *twist^−/−^* and WT tentacles. **(D, D’)** Immunofluorescence staining of mature cnidocytes Basitrichous Haplonema, Microbasic Mastigophore, and Spirocyst with NvNcol-3 (green) in both *twist^−/^* and WT tentacles. **(E-F)** DIC microscopy images of Basitrichous Haplonemas, Microbasic Mastigophores, and Spirocysts found in both WT and *twist^−/−^* mutants **(E’-F’)**. Scale bars are 50µm white and 5µm blue.

**Supplementary Figure 5.**
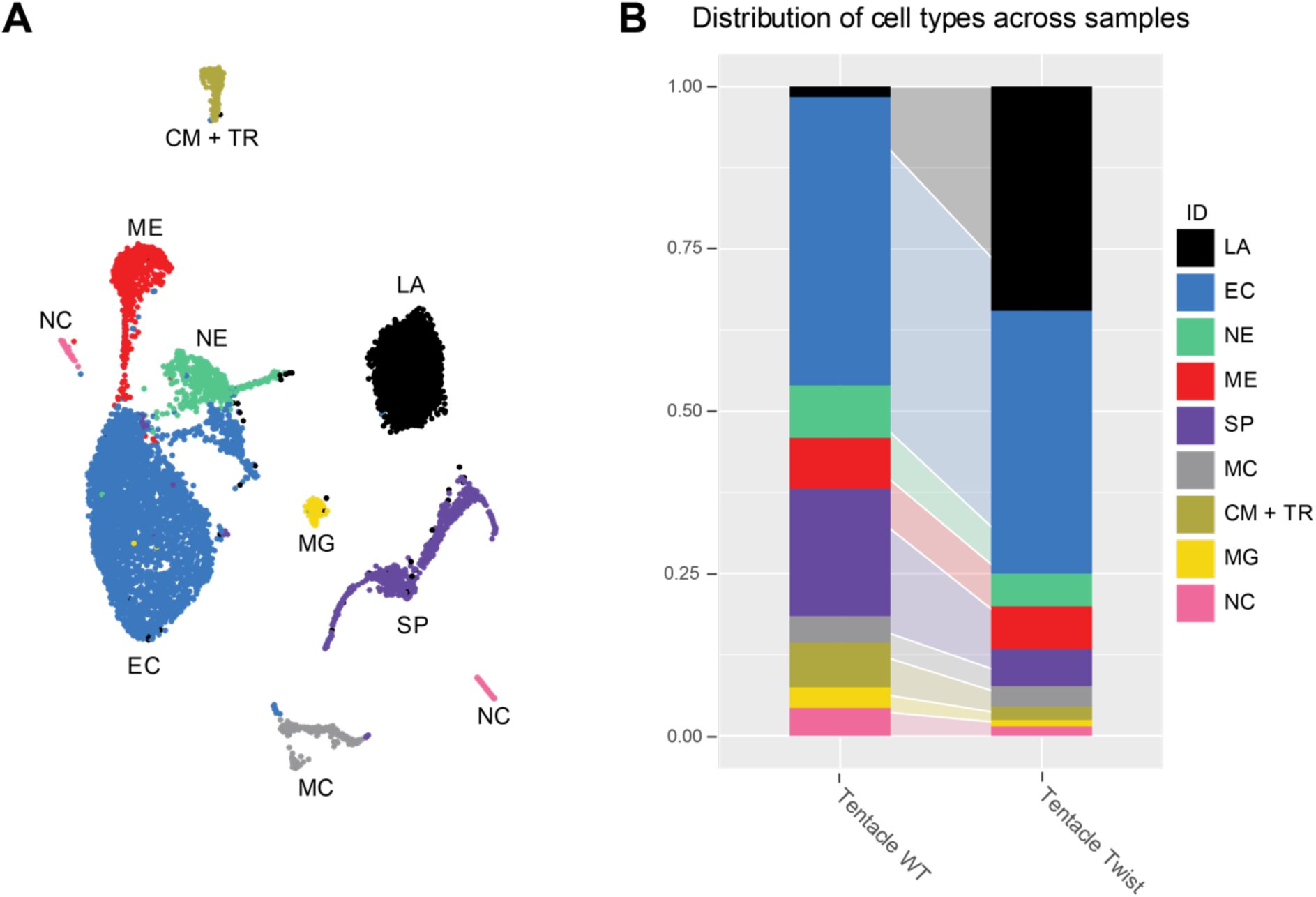
scRNA-seq Analysis of WT and *twist^−/−^* tentacle libraries. **(A)** Dimplot of both scRNA-seq libraries (Tentacle WT and Tentacle *twist^−/−^*) showing the main clusters found in each library. ME: mesoderm, EC: ectoderm, MG: mucus glands, CN: cnidocytes, MC: mature cnidocytes, SP: spirocytes, CR: circular muscle, TR: tentacle retractor muscle, NE: neurons, NC: not categorized, LA: library artifact. **(B)** Cell type distribution across all samples, showing no absence of characterized clusters but lower ratio of neurons, spirocytes and muscle cell populations in the *twist^−/−^* library.

**Supplementary Figure 6.**
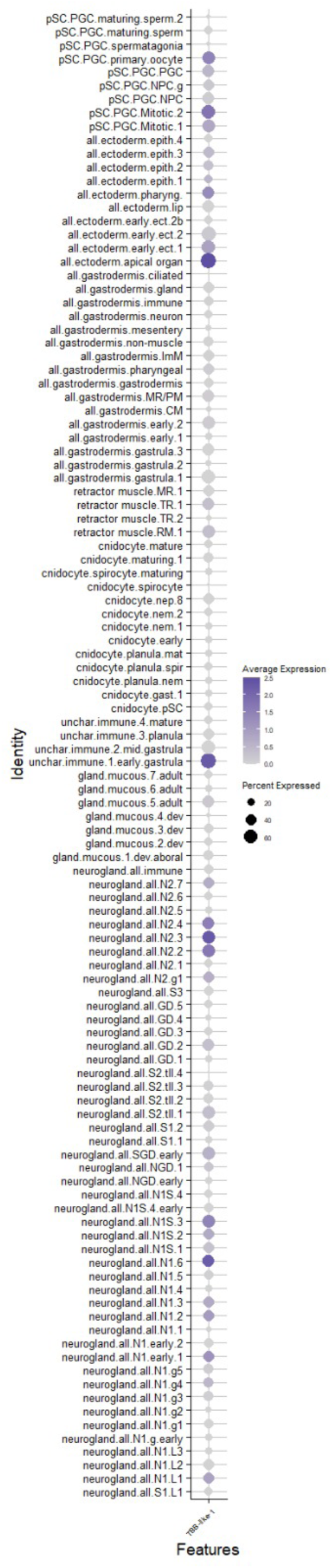
*tbb-like1* is broadly expressed with enrichment in specific neural subtypes. Dot plot showing expression of *tbb-like1* across all annotated cell clusters in the integrated scRNA-seq dataset. Dot size represents the percentage of cells expressing the gene within each cluster; color intensity reflects average expression level. While *tbb-like1* is detected in nearly all clusters, moderate enrichment is observed in a subset of neural subtypes.

